# Robustness and variability in *Caenorhabditis elegans* dauer gene expression

**DOI:** 10.1101/2024.08.15.608164

**Authors:** Johnny Cruz Corchado, Abhishiktha Godthi, Kavinila Selvarasu, Veena Prahlad

## Abstract

Both plasticity and robustness are pervasive features of developmental programs. The dauer in *Caenorhabditis elegans* is an arrested, hypometabolic alternative to the third larval stage of the nematode. Dauers undergo dramatic tissue remodeling and extensive physiological, metabolic, behavioral, and gene expression changes compared to conspecifics that continue development and can be induced by several adverse environments or genetic mutations that act as independent and parallel inputs into the larval developmental program. Therefore, dauer induction is an example of phenotypic plasticity. However, whether gene expression in dauer larvae induced to arrest development by different genetic or environmental triggers is invariant or varies depending on their route into dauer has not been examined. By using RNA-sequencing to characterize gene expression in different types of dauer larvae and computing the variance and concordance within Gene Ontologies (GO) and gene expression networks, we find that the expression patterns within most pathways are strongly correlated between dauer larvae, suggestive of transcriptional robustness. However, gene expression within specific defense pathways, pathways regulating some morphological traits, and several metabolic pathways differ between the dauer larvae. We speculate that the transcriptional robustness of core dauer pathways allows for the buffering of variation in the expression of genes involved in adaptation, allowing the dauers induced by different stimuli to survive in and exploit different niches.

## BACKGROUND

Understanding how genotypes map onto phenotypes remains an exciting and unsolved problem in biology^1^. Phenotypic plasticity, the process by which a single genotype can yield several phenotypes, allows organisms to adapt their development, physiology, metabolism, and even morphology to a varying and unpredictable environment^2,3^ (Fig 1A). However, robustness, the invariant expression of a trait despite genetic or environmental perturbations, is also essential for survival and fitness and is a pervasive feature of developmental programs ^4–6^(Fig 1B).

**Figure 1.**
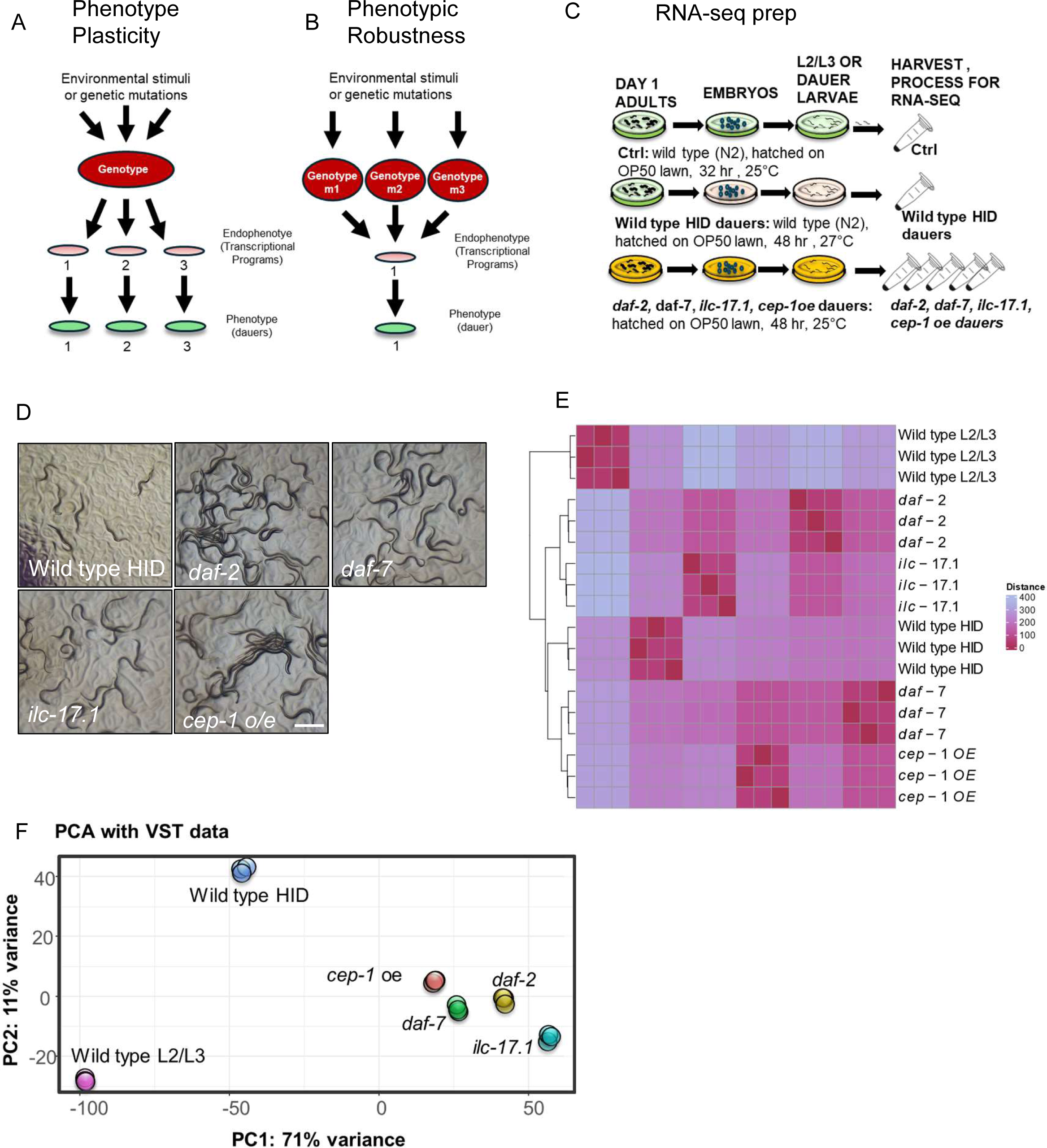
Experimental design. **A, B.** Models for phenotypic plasticity (A) and phenotypic robustness (B). **C.** Schematic of experimental design: RNA sequencing (RNA-seq) of high temperature-induced wild-type dauers (N2; *<u>H</u>*igh *<u>T</u>*emperature *I*nduced *<u>D</u>*auers or wild type HID or WT HID), *daf-2 (e1370) III*, *daf-7(e1372) III* and *ilc-17.1 (syb5296) X* dauers, and dauers that result from overexpressing *cep-1 (cep-1oe).* Samples were collected as described in the text. Wild-type larvae that were grown for 32 hours post-hatching at 25°C to reach the late L2/early L3 stage were used as comparisons. **D.** Micrographs of dauer larvae as used for RNA-seq. Scale bar=1 mm. **E.** Pair-wise distance matrix of RNA-seq samples shows the expected clustering of total RNA of the biological triplicates of each strain [Strains used: wild type (N2) L2/L3, wild type (N2) HID, *daf-2 (e1370) III*, *daf-7(e1372) III* and *ilc-17.1 (syb5296) X* and *cep-1 (cep-1oe)*. **F.** Principal Component Analysis (PCA) of the three repeats of RNA-seq samples.

Dauer formation in the nematode *Caenorhabditis elegans* is considered an example of extreme phenotypic plasticity^7–14^. The *C. elegans* dauer stage is an alternative developmental stage to the third larval stage of the nematode triggered during late larval stage1 (L1)/early larval stage 2 (L2) by environmental stressors such as starvation, crowding, or extreme temperatures. Mutations that downregulate growth-promoting signals that license development also promote dauer arrest and are thought to mimic the environmental triggers^7–14^. Thus, the insulin signaling pathway (ILS) whereby insulins released in the presence of food act through the sole insulin-like receptor, DAF-2, to antagonize the activation of the Forkhead transcription factor DAF-16/FOXO is required for continuous growth; downregulating DAF-2 signaling, as occurs during food scarcity leads to dauer arrest. Likewise, the TGF-β pathway, where the DAF-7 ligand acts through the TGF-β receptors DAF-1/DAF-4 to antagonize the DAF-3/SMAD-DAF-5/Ski transcription factor complex, is also required to permit continuous growth and *daf-7* mutants constitutively arrest as dauers under conditions where wild-type larvae continue development into reproductive adults^15–17^. Recently a cytokine interleukin IL-17 pathway that inhibits the *C. elegans* p53 ortholog p53/CEP-1 has also been shown to be necessary for continuous growth^18^.

Dauer entry is accompanied by a developmental arrest, gene expression changes^19^, morphological changes such as radial shrinkage, pharyngeal constriction, development of a specialized cuticle and buccal plug, physiological changes such as a precipitous decrease in feeding, behavioral changes such as the favoring of nictitation, and metabolic changes such as reduced activities of glycolytic, gluconeogenic, Tricarboxylic Acid Cycle (TCA) cycle, and oxidative phosphorylation pathways^7–10,20^. The different environmental and genetic triggers responsible for these profound changes are thought to act largely through independent or parallel mechanisms to initiate dauer entry^7,8^. This has been well studied in the case of the ILS/DAF-2 and TGF-β/DAF-7 pathways, which regulate parallel and independent inputs into the dauer decision^7–10,20^, although they share some pathway components: insulin gene *daf-28* is regulated by both ILS/DAF-2 and TGF-β/DAF-7 pathways^21^. Similarly, although both loss of DAF-2 and ILC-17.1 acts genetically upstream of DAF-16/FOXO for dauer arrest, they appear to operate through different mechanisms since *ilc-17.1* mutants also require active CEP-1/p53 to trigger arrest, but *daf-2* mutants do not^18^. All dauer pathways impinge on the steroid hormone signaling pathway, DAF-12, but DAF-12 activity has complex effects on dauer arrest as well as developmental timing, and certain *daf-12* mutations cause constitutive dauer arrest while others prevent dauer formation^7–9,11^. In addition, several other mutations and environments modulate these primary triggers to favor or disfavor dauer entry^7–14^. Thus, it is unclear whether gene expression in the different dauer larvae induced to enter dauer by different environmental or genetic perturbations varies based on the route into dauer or is invariant and robust, independent of the genetic or environmental trigger that induces dauer arrest.

To address this question, we used next-generation RNA sequencing (RNA-seq) to obtain the transcriptional profiles of different dauer larvae where developmental arrest was triggered by different genetic or environmental stimuli: dauers initiated by exposure to high temperatures (N2; *<u>H</u>*igh *<u>T</u>*emperature *I*nduced *<u>D</u>*auers or HID)^22^, loss of the insulin signaling (*daf-*2)^23^, downregulated TGF-β signaling (*daf-*7)^24^, loss of the cytokine ILC-17.1 pathway (*ilc-17.*1), and activation of the CEP-1/p53 pathway (*cep-1 o/*e)^18^. We then compared the transcriptomes of the dauer larvae. We assessed the extent to which gene expression varied between these dauers by computing the coefficient of variation (CV; standard deviation divided by the mean) of all expressed genes across all the dauer larvae. Separately, we evaluated concordance in gene expression by measuring Spearman’s rank-order correlation between the expression levels of genes within different Gene Ontologies (GO) and functional pathways, pair-wise in all dauer larvae, to compare overall patterns of gene expression. Our analysis shows that while the expression levels of most genes vary widely between the different dauer larvae, expression patterns within the majority of functional pathways are strongly correlated, implying the presence of robust constraints that stabilize gene expression. However, specific stress, defense, and metabolic pathways were poorly correlated between almost all dauer larvae. We speculate that the robustness of core dauer pathways buffers the variability in gene expression in specific pathways that modulate adaptation to the environment, allowing the different dauer larvae to better exploit and survive in different environmental niches.

## RESULTS

### Dauers induced by different environmental and physiological stimuli utilize similar processes for dauer arrest

To assess the gene expression profiles of dauer larvae generated under different conditions, we conducted RNA sequencing (RNA-seq) of high temperature-induced wild-type dauers (N2; *<u>H</u>*igh *<u>T</u>*emperature *I*nduced *<u>D</u>*auers or wild-type HID; WT HID), *daf-2 (e1370) III*, *daf-7(e1372) III* and *ilc-17.1 (syb5296) X* dauers, and dauers that result from overexpressing the *C. elegans* p53-like gene *cep-1 (cep-1oe)* (Fig. 1C, D). Wild-type HID were generated by allowing wild-type larvae to grow at 27°C for 48 hours following hatching. Dauers from the *daf-2 (e1370) III*, *daf-7(e1372) III* and *ilc-17.1 (syb5296) X* and *cep-1 oe* backgrounds were generated by allowing embryos to grow at 25°C for 48 hours following hatching. We confirmed that all dauer larvae displayed the characteristic morphological features of dauer (Fig 1D). In addition, to ensure that the conditions used generated ‘true’ dauers, we tested their resistance to 1% SDS in a separate pilot experiment, and verified that the larvae generated in response to these conditions were SDS-resistant^8^. The <1% larvae that escaped dauer arrest were visible as stage 4 larvae (L4) at the time of harvesting and were manually removed prior to mRNA extraction. Thus, the mRNA was highly enriched for dauer-specific genes expressed upon arrest, 48 hours post-hatching. To identify genes that were differentially expressed (differentially expressed genes; DEGs) during dauer arrest in each of the dauer types, we compared their RNA with RNA sequenced from late L2/early L3 wild-type larvae that were grown for 32 hours post-hatching at 25°C. Three independent biological samples were sequenced. The L2/L3 larvae are considered to be at a comparable developmental stage to dauer but do not arrest as dauers and instead are fated to develop into reproductive adults. Their growth at 25°C allowed us to account for the effects of temperature on development.

A sample distance matrix produced by comparing expression levels across all genes between samples demonstrated excellent agreement among biological replicates (Fig. 1E; Supplementary Table 1). The mean expression levels (log_10_ TPM) of all genes were comparable but were modestly lower in the *daf-7* and *ilc-17.1* larvae, perhaps suggestive of global differences in transcription (Supplementary Fig. 1A). Principal Component Analysis (PCA) showed that all dauers separated well from continuously growing larvae, and clustered according to biological replicates (Fig. 1F). Importantly, the different dauer types also separated from each other, indicating that they differed from each other, with the separation of wild-type HID from other dauers along PC1 and PC2 being larger than the more modest separation between the *daf-2, daf-7*, *ilc-17.1 or cep-1oe* dauers (Fig. 1F).

As a first step towards evaluating similarities and differences between the dauer larvae, we identified the genes that are differentially expressed (differentially expressed genes; DEGs) in each of the dauers, by comparing their gene expression to that in continuously developing L2/L3 larvae and conducted Gene Ontology (GO) enrichment analysis^25–29^. The different dauers displayed similar numbers of DEGs (N2 HID =16302, *daf-2* dauers=16885, *daf-7* dauers=16557, *ilc-17.1* dauers=16747 and *cep-1oe* dauers*=* 16565, padj< 0.05), of which the majority (13120 DEGs) were common (Fig 2A; Supplementary Table 2). The DEGs in the different dauers largely were enriched in similar processes, arguably those necessary for developmental arrest (padj< 0.05; Fig. 2B, C; Supplementary Figs 2 and 3). Thus, all five types of dauer larvae downregulated genes regulating the cell cycle, DNA metabolic processes, translation, and other anabolic processes. Genes contributing to neuropeptide signaling and ion homeostasis were upregulated in all dauers. In addition to these similarities, there were some differences: some GO pathways were enriched in some dauers, but not others (Fig. 2B-C; Supplementary Figs 2 and 3; Supplementary Tables 4 and 5), and a small group of between 27 to 109 genes were uniquely altered in only one of the dauer larvae (Fig 2A; Fig 2D; Supplementary Table 3). For instance, *daf-2* dauers differed from other dauers in the expression of genes related to striated muscle-dense body and contractile fiber and uniquely altered the expression of genes enriched in “molecular transducer activity” (*srb-6, str-220*, *srg-2, etc.*; Fig 2D; Supplementary Table 6). Similarly, DEGs in larvae that were induced to arrest as dauers due to the loss of the interleukin cytokine gene *ilc-17.1* were not enriched for several immune response categories that were enriched in all other dauer larvae but had an over 20-fold enrichment in GO categories ‘ciliary plasm’ (downregulation of *klp-11* and upregulation of H13N06.7) and ‘sodium channel activity’; Fig. 2D; Supplementary Table 6). Likewise, DEGs in *daf-7* dauers also differed in enrichment for a smaller subset of “innate immune response” genes. The GO categories ‘postsynaptic membrane’, ‘regulation of postsynaptic membrane potential’ and “stabilization of membrane potential” were unique to DEGs in the *daf-*7 dauers, consistent with what is known regarding DAF-7 expression in neurons, and its effects on the *C. elegans* nervous system and behavior^30,31^ (Fig. 2D; Supplementary Table 6). These data suggested that the different dauer larvae likely utilized similar processes for dauer arrest and other physiological functions, but also displayed differences in gene expression.

**Figure 2.**
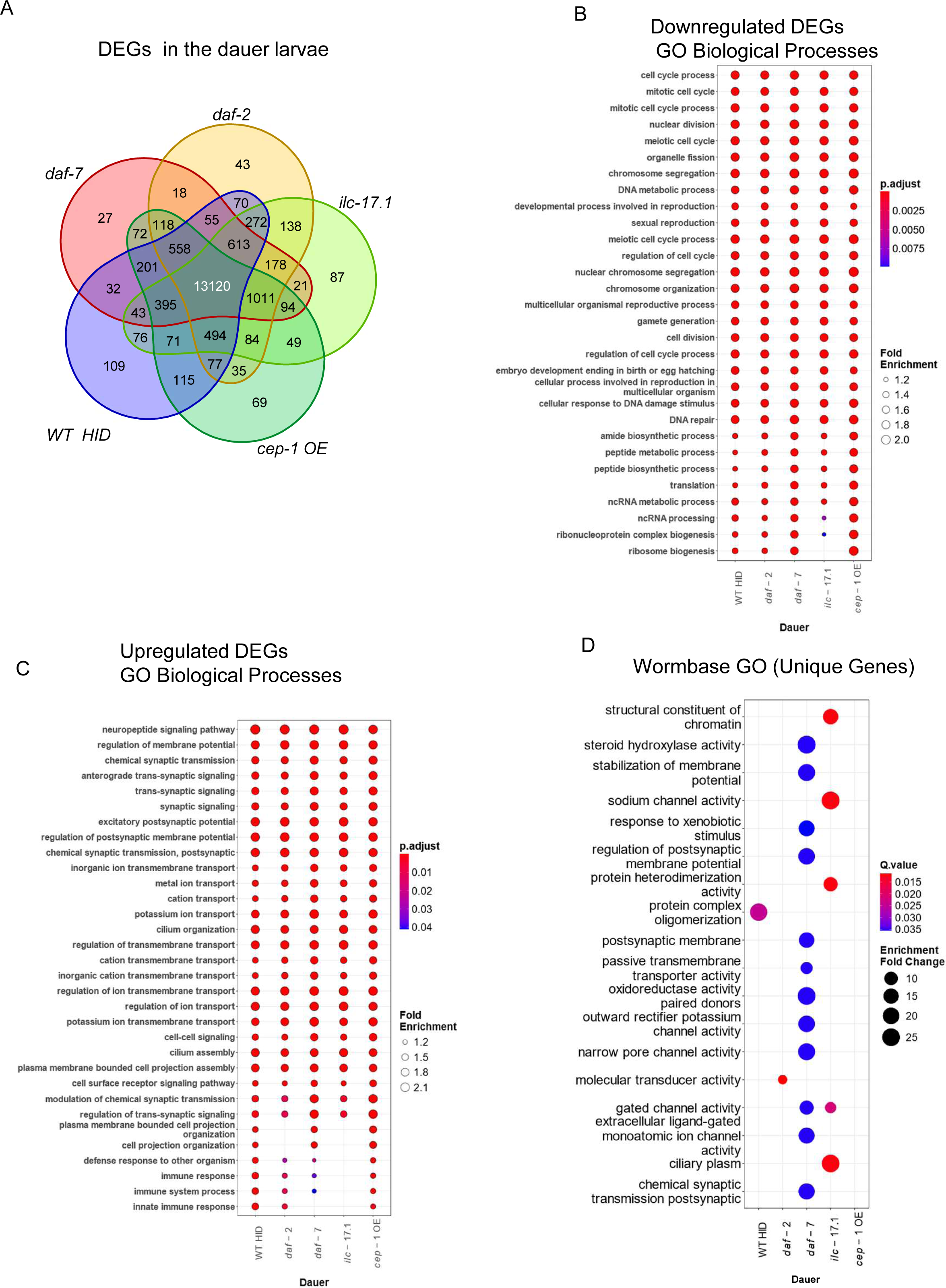
Dauers induced by different environmental and physiological stimuli utilize similar processes for dauer arrest. **A.** Venn Diagram showing overlap between differentially expressed genes (DEGs) in wild-type HID, *daf-2, daf-7*, *ilc-17.1* and *cep-1oe* dauers compared to wild type L2/L3 larvae. **B.** Dot plot showing comparison of enrichment between downregulated DEGs in wild-type HID, *daf-2, daf-7*, *ilc-17.1* and *cep-1oe* dauers compared to wild type L2/L3 larvae. Y axis: GO categories (Biological Processes). Color bar: adjusted p-values (Benjamini-Hochberg corrected, p<0.05), lower p-value in red, higher p-value blue. Circle size: Fold Enrichment (Gene Ratio/Background Ratio). **C.** Dot plot showing comparison of enrichment between upregulated DEGs in wild type HID, *daf-2, daf-7*, *ilc-17.1 or cep-1oe* dauers. Y axis: GO categories (Biological Processes). Color bar: adjusted p-values (Benjamini-Hochberg corrected, p<0.05), lower p-value in red, higher p-value blue. Circle size: Fold Enrichment. **D.** Dot plot showing enrichment of DEGs unique to wild type HID, *daf-2, daf-7*, *ilc-17.1 or cep-1oe* dauers. Y axis: GO categories (Wormbase). Color bar: adjusted p-values (Q.value<0.05), lower p-value in red, higher p-value in blue. Circle size: Fold Enrichment.

### Different dauers display high variability in gene expression levels but strong correlations in gene expression patterns

To evaluate the extent to which gene expression in the five dauer larvae was similar or differed, we (i) estimated the variance in expression of all expressed genes (mean expression>10 tpm), using the coefficient of variation (CV) as an indicator of variance^32–34^, and (ii) computed the Spearman’s rank-order correlation, *rho*, between all genes in pairs of dauer larvae as a measure of similarity in the patterns or relationships between genes. The CV was computed for each gene by dividing the standard deviation (SD) of its expression across all dauers by its mean expression, and ranged from 11% to 387%. Strikingly, over half the genes (8538/16940, 50.4%) exhibited high CVs of >50%, where the SD values were over half the mean expression values (based on the genome-wide CV distribution, we defined CV < 30% as low, <30% < CV < 50% as moderate, and CV > 50% as high; Fig 3A; Supplementary Table 7). The high CVs were not simply a consequence of low gene expression, as seen by plotting the SD as a function of the mean and inferred because the CVs were computed after filtering for low mean expression (Supplementary Fig 1B; Materials and Methods). This suggested that notwithstanding the enrichment of similar processes by the upregulated or downregulated genes of all dauer larvae, the expression levels of genes contributing to these processes varied, and in some cases varied widely, across the different dauers.

**Figure 3.**
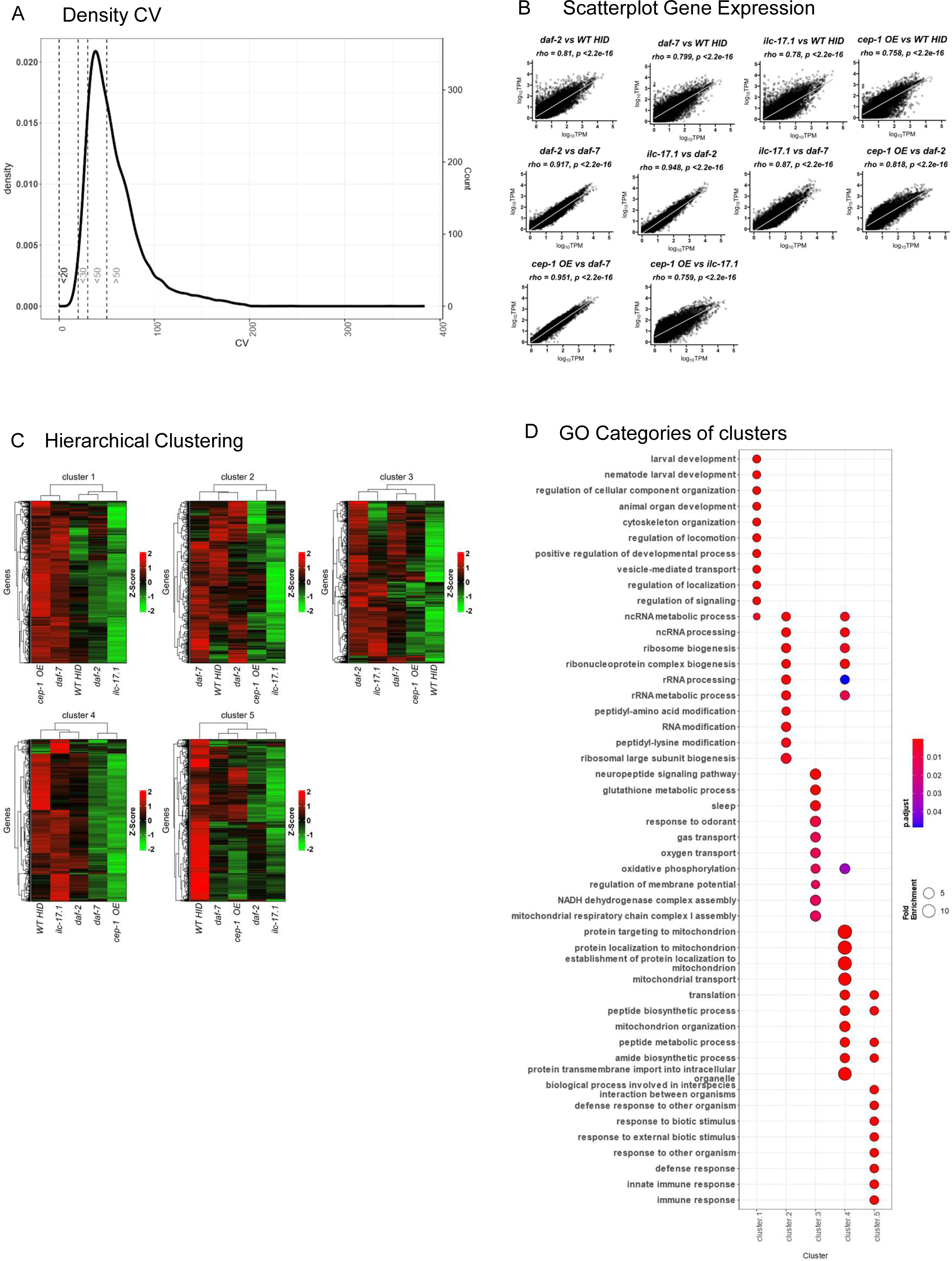
Dauers display high variability in gene expression levels but strong correlation in gene expression patterns. **A.** Density plot of the Coefficient of Variation (CV). X-axis: CVs of all expressed genes (mean expression >10 counts) computed by dividing the standard deviation (SD) across all dauers by its mean expression across all dauers. Dotted lines demarcating CVs: low, (CV < 30), moderate (30 < CV < 50), high (CV > 50). Y-axis (left) density, (right): gene count. **B.** Scatter plots showing pairwise Spearman correlation between the wild type (N2) HID, *daf-2 (e1370) III*, *daf-7(e1372) III* and *ilc-17.1 (syb5296) X* and *cep-1 (cep-1oe)*. Line represents linear regression. TOP: Dauers that are compared, and Spearman rho values shown. p-value is corrected for multiple tests; Benjamini-Hochberg. **C.** Heatmap depicting relative gene expression (Log_10_TPM) as Z-scores across the dauers. The five different clusters were obtained by hierarchical clustering (method: complete, distance: Euclidean) between the genes and applying the elbow method to select the optimal number of clusters. TOP: Cluster number. X-axis: dauers. Y-axis: genes. **NOTE: Dauer order on X-axis differs in each cluster.** Columns are ordered based on the Hierarchical clustering (method: complete, distance: Euclidean) between the different dauers, as shown in the dendrogram on top of the Heatmap. Green-Black-Red Color bar: Z-scores. **D.** Dot plot comparison of enrichment between of five clusters identified in B. Y-axis: the GO categories. The color bar shows adjusted p-values (Benjamini-Hochberg corrected, p<0.05), lower p-values in red, and higher p-values in blue. Circle size: Fold Enrichment.

Surprisingly, Spearman’s rank-order correlation tests between pairs of dauers showed the opposite trend to what was expected from the large dispersion in gene expression levels, and gene expression patterns in all the different dauers were strongly correlated (Fig 3B). Thus, the Spearman’s correlation coefficient, *rho*, calculated between the log_10_ gene expression values of expressed transcripts in the five dauer larvae ranged between >0.7, to >0.9 [we defined rho=0.7-1.0 as a strong correlation, rho=0.5-0.7 as moderate, and rho<0.5-0.6 as weak correlation, padj<0.05, according to ^1,12^]. The correlation coefficient between the N2 HID dauers and *daf-2, daf-7*, *ilc-17.1 or cep-1oe* dauer larvae was rho(16085)=0.81, rho(16085)=0.799, rho(16085)=0.78, and rho(16085)=0.758 respectively (p=2.2e-16 in all cases). Surprisingly, gene expression in *daf-2* and *daf-7* dauers were also concordant [rho(16085) =0.917], even though *daf-2* and *daf-7* mutations activated separate, independent mechanisms to trigger dauer entry. The more recently discovered dauer pathway triggered by the loss of *ilc-17.1*, which decreases glucose uptake, resembled *daf-2* dauers, which downregulate insulin signaling [rho(16085) =0.948] but differed more from *daf-7* dauers. Dauers induced by *cep-1* oe most resembled *daf-7* dauers and most differed from *ilc-17.1* dauers (rho(16085) =0.951 and rho(16085) =0.759 respectively).

Hierarchical clustering of gene expression (log_10_ TPM) also supported the existence of similarities and differences in gene expression between the different dauers. Genes expressed in all dauer larvae separated into five clusters (Supplementary Fig. 1D), and different subsets of genes were similar in different subsets of dauer larvae (Fig. 3C, D; Supplementary Tables 8, 9). For instance, *daf-7* and *cep-*1 *oe* dauers were similar and distinctly different from the other dauers in their expression of genes in Cluster 1 and Cluster 4 (Fig 3C; Supplementary Tables 8,9), which were enriched in GO categories that regulated different aspects of larval development and locomotion and mitochondrial protein targeting, organization, and transport respectively (Fig 3D). Likewise, *daf-*2, *daf-*7, and wild-type HID were more similar to each other and different from *ilc-17.1* and *cep-1 oe* dauers in the expression of genes related to RNA processing (cluster 2), all dauers separated from wild-type HID with regards to neuropeptide signaling, several behavioral responses (cluster 1), and protein translation, peptide biosynthetic processes and defense and immune programs (cluster 5) (Fig. 3C, D; Supplementary Tables 8, 9).

Thus, together, these data showed that gene expression levels varied between the different dauers, with over 50% of all expressed genes displaying CVs of >50%. Hierarchical clustering of gene expression showed that different subsets of genes enriched in different biological pathways were similarly expressed or differed from each other in different subsets of dauers, likely driving the high CVs. However, surprisingly, the relative patterns of gene expression appeared to be preserved in the different dauer larvae, as indicated by the strong Spearman’s rank-order correlation coefficients in all the pair-wise comparisons.

### Gene expression in the core dauer pathway is robust

These observations suggested that gene expression patterns were concordant within pathways despite the high variation in expression levels of the individual genes involved in these pathways. We tested whether this was the case by examining seven ‘core’ dauer pathways. As ‘core dauer pathways’, we chose the ILS (insulins and DAF-16 targets), TGF-β and DAF-12/steroid hormone pathways that trigger dauer and are thought to coordinate the organism-wide developmental pause, pathways that implement the growth arrest, namely those that regulate the cell cycle (cyclins, the Anaphase Promoting Complex, cyclin-dependent kinases, cyclin inhibitors, etc.), and energy metabolism (glycolysis and electron transport chain, ETC), and six cuticulin genes that are required for the formation of dauer specific alae^8,9^. The *daf-2* and *daf-7* dauers harbor mutations in the sole insulin receptor and the DAF-7/TGF-β ligand, respectively, and are expected to downregulate signaling through these pathways, whereas the other dauers, i.e., wild-type HID, *ilc-17.1* and *cep-1 oe* dauers are wild-type for genes in these pathways. Therefore, we expected the expression levels of ILS and TGF-β pathway genes, and perhaps also the DAF-12 pathway genes, to vary between all dauers, resulting in high CVs. Thus, if the Spearman’s correlation between gene expression in the ILS and TGF-β pathways (and DAF-12 pathway) of *daf-*2 and *daf-7* dauers when compared to the other dauers remained strong, this could suggest the presence of additional constraints over gene expression that maintained transcriptional robustness.

As expected, the expression levels of ILS (Insulins and DAF-16), and DAF-12 pathway genes differed between the different dauers: the majority of insulins (28/31, 90.32%), DAF-16 targets (56/78, 71.79%) and DAF-12 targets (16/29, 55.17%) had high CVs of >50% (Fig 4A; Supplementary Table 10). Surprisingly, the CVs of the TGF-β genes (20/26, 76.9%) were low to moderate (20%-50%; Fig 4A), even though *daf-7* dauers had a mutation in the DAF-7/ TGF-β ligand and downregulate TGF-β signaling, suggesting that more needed to be understood regarding the transcriptional regulation of this pathway. Nevertheless, despite the variability and high CVs of the genes functioning in the ILS, DAF-16, and DAF-12 pathways, the pair-wise correlation coefficients for these genes computed between all dauer larvae, even between *daf-2* dauers and dauers that were wild-type for the ILS genes, were overwhelmingly strong and significant [rho >0.7 in all cases, and rho>= 0.9 for the majority of comparisons; padj<0.05 (Fig 4B; Supplementary Tables 11)]. As could be expected from the low variability in the expression of TGF-β pathway genes, the same was true for the correlation between genes in the TGF-β pathway between *daf-7* and other dauers, although TGF-β pathway gene expression compared between *ilc-17.1* and *cep-1 oe* dauers, neither of which harbored mutations in TGF-β components, was significant but only moderately correlated [(rho=0.59; padj<0.05); (Fig 4B; Supplementary Tables 11)].

**Figure 4.**
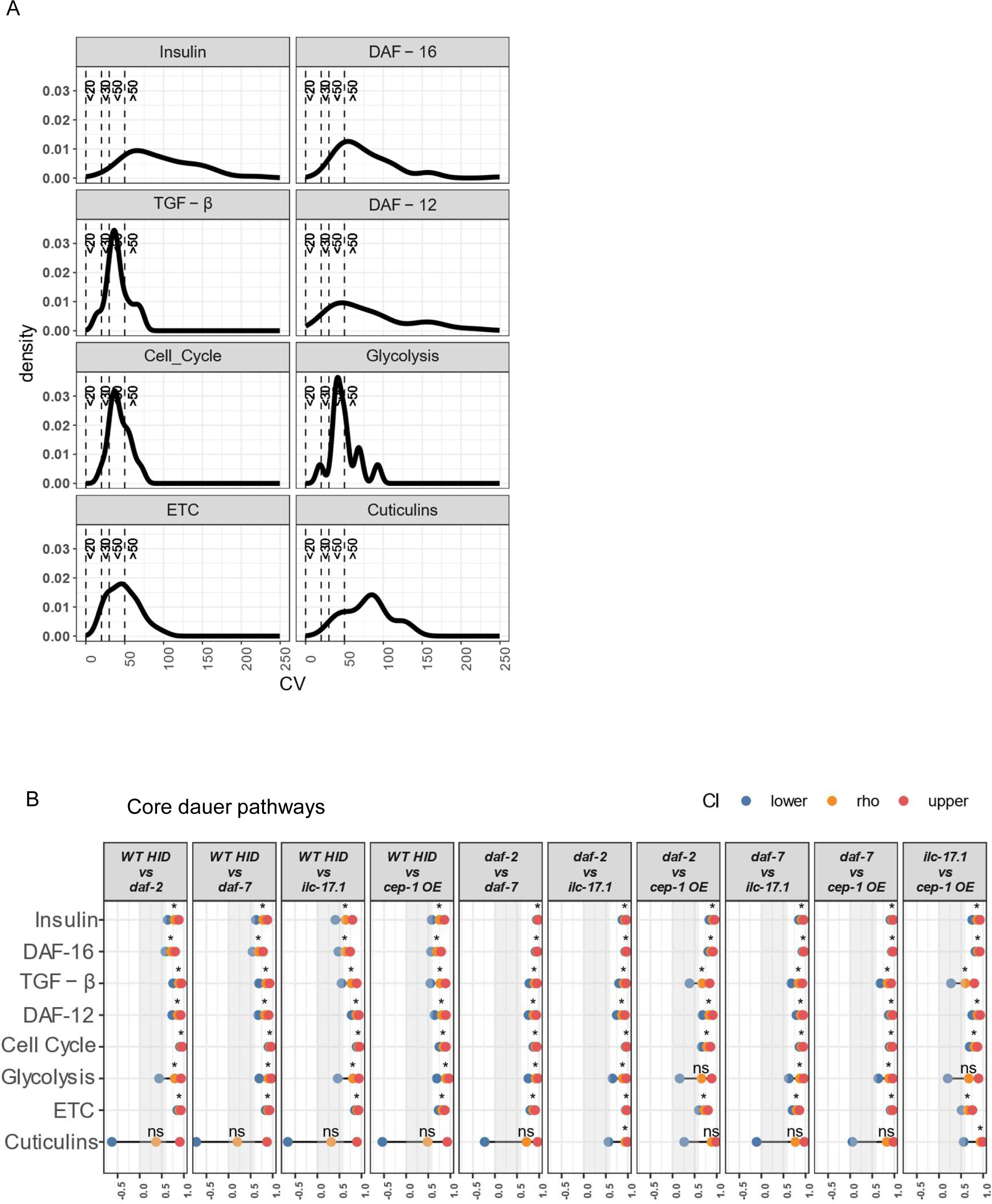
Gene expression in the core dauer pathway is robust. **A.** Density plot of the Coefficient of Variation (CV) of genes in the core dauer pathway. Labels of the pathway on top. X-axis: CVs of all genes in the pathway computed by dividing the standard deviation (SD) of its expression across all dauers by its mean expression across all dauers. Dotted lines demarcating CVs: low, (CV < 30), moderate (30 < CV < 50), high (CV > 50). Y-axis (left) density. **B.** Dumbbell Plot showing Spearman’s correlation coefficient (rho) between pairs of dauers in ‘core’ dauer pathways. TOP labels: the pair of dauers compared. Y-axis: pathway. X-axis: rho value and confidence interval (CI): blue dot represents lower CI, yellow dot represents rho value, and red dot represents high CI. p-values, Bonferroni corrected. *p-value <0.05; ns, not-significant.

Genes that function in cell cycle or energy metabolism pathways were less variable in their expression levels, and the majority of cell cycle (45/67, 67%), glycolysis (8/13, 61.5%), and ETC (49/88, 55.68%) genes had CVs that ranged between moderate to low (20%-50%) and in the lower half of the genome-wide CV range when computed across the different dauers (Fig 4A; Supplementary Tables 10). Accordingly, the correlation between genes expressed in these pathways was strong for all pair-wise comparisons of dauers, with the exception of glycolysis genes compared between *cep-1*oe dauers and *daf-2* or *ilc-17.1* dauers for reasons we cannot explain (Fig 4B; Supplementary Tables 11).

Expression levels of the six cuticulin genes were variable (CV> 50%; Fig 4A), but in contrast to the genes in the ILS and DAF-12 pathways, the expression patterns of these genes were also variable in the different dauer larvae, and the correlation coefficient in most cases was not significant (Figure 4B, Supplementary Tables 10,11). A comparison of the mean mRNA expression values (rlog counts), and the log_2_ fold changes of genes in the core dauer pathway reinforced this interpretation (Supplementary Fig 4A-D).

These data indicate that variance in gene expression can be accommodated or stabilized in specific pathways to yield similar relationships between gene expression, indicative of transcriptional robustness^35^. These data also implicate different mechanisms of control over gene expression in the core dauer pathway to maintain strongly concordant patterns of expression. Thus, the cell cycle and energy metabolism genes that can be expected to implement the dauer arrest appear to be inherently more reliable in their expression, and levels were more similar across the different dauer larvae; correspondingly, their expression patterns in all dauer larvae were also similar. The same was true of the TGF-β pathway. The expression levels of genes in the ILS/ /DAF-12 pathways that trigger the dauer decision were more variable: however, despite their variability in expression levels, the relationship between genes in these pathways was invariant and strongly correlated in all the different dauer types. This was not true for the cuticulin genes which varied in their expression levels and their relative rank ordering.

### Gene expression patterns enriched for a limited set of traits, stress response, immune response, and metabolic pathways vary between the different dauer larvae

Our analysis showed that the different dauer larvae shared and differed in gene expression within specific biological pathways. Therefore, to identify pathways that might distinguish the *daf-2,* daf-7, *lc-17.1,* and *cep-1 oe* dauers, we took advantage of GO enrichment analysis of the DEGs in the different dauer larvae and examined the correlation of gene expression within these GO categories. As shown previously (Fig 2A), a core set of 13120 genes were differentially expressed in all dauers when compared to continuously growing L2/L3 larvae. These shared 13120 DEGs were enriched in 309 GO categories (Fig 5A shows summarized GO categories; Supplementary Table 12 and 13; log_2_ fold change in Supplementary Fig.1C). Gene expression within most of these 309 GO categories was strongly correlated between all the dauers, with the exception of 21 GO categories where correlation was low or not significant (rho <0.6 and/or padj>0.05) between one or more pairs of dauer larvae (Fig. 5B; purple indicated GO categories with low correlation). These weakly correlated GO categories were enriched in genes that acted in several different processes, such as “response to odorants,” “microtubule depolymerization,” “mRNA destabilization,” etc., and varied between specific dauer larvae, and to different extents (Fig 5C). For instance, *daf-2* and *ilc-17.1* dauers differed from wild-type HID and *cep-1 oe* dauers in the expression of genes involved in the “response to odorants”. This GO category (Supplementary Tables 12-14) includes *hsf-1, tph-1,* and nine paralogs of ODR-2 called *hot* genes (for “homologs of odr two”) that have not been widely studied. *ilc-17.1* differed from all other dauers except *daf-2,* and *cep-1 oe* dauers differed from all other dauers besides *daf-7* in the expression patterns of 15 genes that regulate “microtubule depolymerization”, and *cep-1 oe*, dauers differed from all other dauers except *daf-7* in “nuclear-transcribed mRNA catabolic process” and “mRNA destabilization” (Fig. 5C; Supplementary Tables 12-14). Intriguingly *cep-1 oe* dauers also differed from *ilc-17.1* and *daf-2* dauers in their expression of genes regulating dendrite development in GO categories “dendrite morphogenesis” and “regulation of dendrite morphogenesis”, and *ilc-17.1* dauers differed from wild type HID and *cep-1* oe dauers in several aspects of male morphogenesis such as “nematode male tail mating organ morphogenesis”, “male genitalia development”, and “male genitalia morphogenesis” (Fig. 5C; Supplementary Tables 12-14).

**Figure 5.**
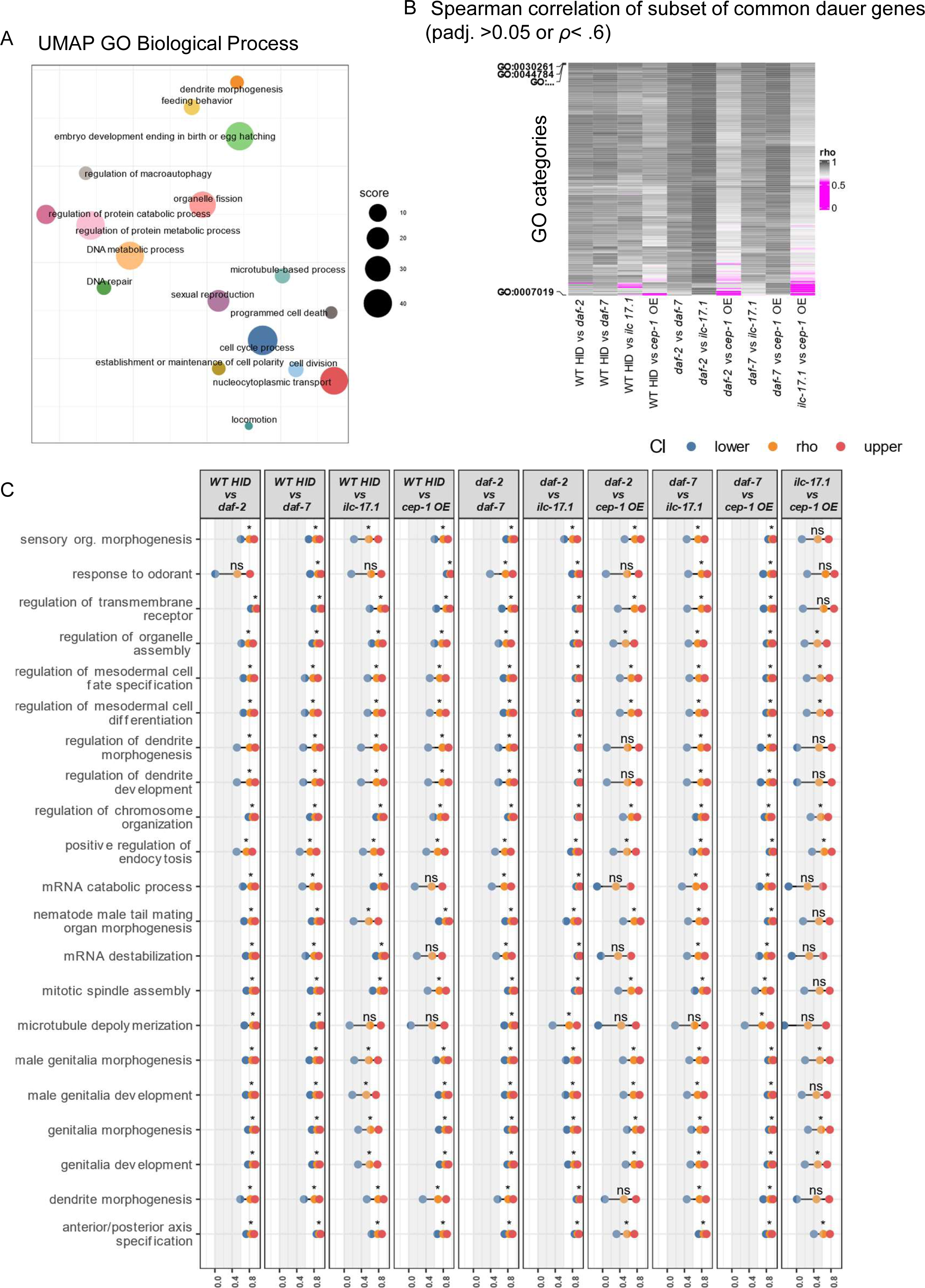
Correlation within GO categories enriched by genes differentially expressed by all dauers shows variation in a small subset of categories. **A.** Scatter plot showing summarized GO categories (Biological Process) enriched from 13210 differentially expressed genes (DEGs) common to all dauers [dauers compared to wild type L2/L3 larvae]. Axes are represented by two UMAP components. Distance between points represents similarity between terms. Size of the points: score in the dissimilarity matrix. **B.** Heatmap depicting Spearman’s correlation coefficient (rho) for pairwise comparisons between all dauers in each GO category enriched, as in A. Colorbar: Black-white: rho >0.6, purple: rho<0.6 or padj Bonferroni corrected >0.05 (indicates low correlation). **C.** Dumbbell Plot showing Spearman’s correlation coefficient (rho) between ‘common dauer genes’ compared between pairs of dauers and analyzed within GO categories depicted in A. TOP labels: the pair of dauers compared. Y-axis: GO categories that were collapsed in A. X-axis: rho values and confidence interval (CI): blue dot represents lower CI, yellow dot represents rho value, and red dot depicts high CI. p-values, Bonferroni corrected. *p-value <0.05; ns, not-significant.

These results suggested that despite the robustness in overall gene expression, the different dauer larvae that arrested development in response to different triggers or stimuli, may also differ in specific physiological and morphological traits.

We next examined whether the different dauer larvae differed in the expression of genes that are co-expressed to constitute the thirty-four gene expression ‘mountains’, as defined by Kim et al (2001)^36^. These gene expression maps or ‘mountains’ were identified by analyzing the co-expression of 5361 *C. elegans* genes across a large number of individual experiments that used wild-type and mutant animals at different developmental stages. The ‘mountains’ fall into categories such as ‘protein expression,’ ‘histone,’ ‘mitochondria,’ ‘germline,’ ‘G protein receptors’, ‘heat shock,’ etc., likely representing genes that act together in a functional and/or spatially coordinated manner. As with the previous analysis, gene expression of all the Kim ‘mountains’ with only six exceptions were also highly correlated between all the dauers (*rho* values were > 0.7 for all ‘mountains’ except those purple areas in the heatmap; Fig. 6A, B; Supplementary Tables 15 and 16). The six exceptions, “heat shock”, “cytochrome p450 (CYP)”, “retinoblastoma complex”, “mechanosensation, “intestine,” and “amine oxidases,” varied either between all dauers or between specific dauer larvae (Fig. 6 A, B; Supplementary Fig. 6 and 7; Supplementary Tables 15 and 16). Thus, the expression of the amine oxidases, which in *C. elegans* consist of a family of seven genes, did not correlate between any two dauer larvae (padj=0.1-1 for all comparisons; Fig. 6A, B). Amine oxidases are heterogenous enzymes that catalyze the oxidative deamination of polyamines and play essential roles in several biological processes in all organisms^37–39^. In *C. elegans,* the putative amine oxidase, *amx-2* is thought to be the essential monoamine oxidase in serotonin and dopamine degradation and is important for lifespan regulation and modulation of the RAS/MAPK pathway activity during *C. elegans* vulval development^40^. The putative monoamine oxidase *gdh-1* is thought to be a mitochondrial gene that plays a role in the innate immune response and susceptibility of *C. elegans* to *Pseudomonas aeruginosa* PAO1 pathogenesis^41^, and the putative monoamine oxidase *hpo-15* (hypersensitive to POre-forming toxin 15)^42^, has also been implicated in a role in defense response to bacterial pathogens. Consistent with the role of overarching role of the p53 family of genes in the control of the cell cycle in all animals, *cep-1* oe dauers differed from all other dauers in the expression pattern of their retinoblastoma complex genes (Fig. 6 A, B; Supplementary Fig. 6B). Intriguingly, amongst the retinoblastoma complex, the log_2_ gene expression changes for *mdt-22* varied to a larger extent in *daf-7* and *cep-1 oe* larvae. *mdt-22* is one of several mediator complex subunits that collaborates with CKI-1, a member of the p27 family of cyclin-dependent kinase inhibitors (CKIs) which themselves are a target of CEP-1/p53, to maintain cell cycle quiescence of vulval precursor cells during larval development^43^. CYP gene expression varied between wild-type HID dauers and *ilc-17.1* dauers, that in the ‘heat shock’ mountain, which consisted mainly of the molecular chaperones, varied between *cep-1* of dauers and *daf-2* dauers (Fig 6 A, B; Supplementary Fig. 7), *cep-1* of dauers and daf-2 dauers also differed from each other in their expression of specific intestine-related genes and in the genes that affect mechanosensation.

**Figure 6.**
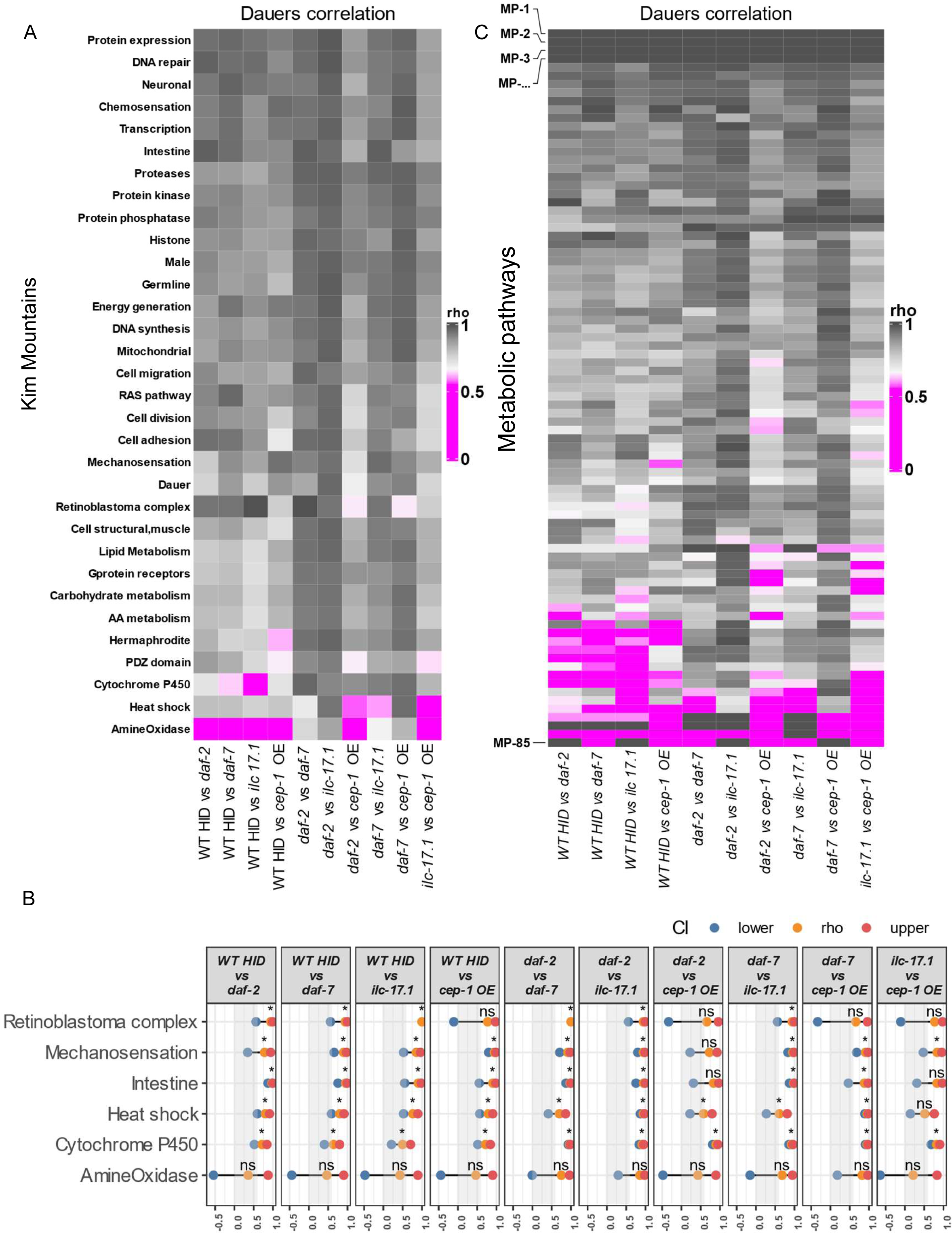
Correlation within categories of co-expressed genes in the *C. elegans* gene expression map [Kim et al. (2001)] and the iCEL1314 metabolic network [Nanda et al. (2023)] shows that dauers vary in stress and metabolic pathways. **A.** Heatmap depicting Spearman’s correlation coefficient (rho) for pairwise comparisons between all dauers in Kim ‘mountains’ from the *C. elegans* gene expression map [Kim et al (2001)]. Colorbar: Black-white: rho >0.6, purple: rho<0.6 or padj Bonferroni corrected >0.05 (indicates low correlation). **B.** Dumbbell Plot showing Spearman’s correlation coefficient (rho) between pairs of dauers compared within ‘Kim mountains’. TOP labels: the pair of dauers compared. Y-axis: pathway. X-axis: rho value and confidence interval (CI): blue dot represents lower CI, yellow dot represents rho value, and red dot depicts high CI. p-values, Bonferroni corrected. *p-value <0.05; ns, not-significant. **C.** Heatmap depicting Spearman’s correlation coefficient (rho) for pairwise comparisons between all dauers within metabolic pathways in the iCEL1314 metabolic network [Nanda et al. (2023)]. Colorbar: Black-white: rho >0.6, purple: rho<0.6 or padj Bonferroni corrected >0.05 (indicates low correlation).

Finally, we examined correlation in the expression of 1,799 genes within 85 metabolic pathways described in the iCEL1314 metabolic network model of Nanda et al. (2023)^44^, where the genes also clustered based on their co-expression in metabolic pathways. In contrast to the overarching high correlation seen across the GO pathways and the Kim mountains, where only a limited number of pathways differed between the different dauer larvae, gene expression in over a third of the iCEL1314 metabolic pathways (thirty-four of the 85 pathways) showed variability between two or more dauer larvae (we discarded three pathways since they only contained two genes each; Fig 6C, Fig 7A; Supplementary Tables 17 and 18). Thus, gene expression in the glycine cleavage pathway, histidine degradation, mevalonate metabolism, and propionate degradation pathways differed between all dauer larvae, and did not correlate significantly between practically any pair of dauer larvae [padj>0.05; (Fig 7A; Supplementary Figs 8-11). Wild-type HID differed from all other dauers in the expression of genes involved in ROS metabolism, folate cycle, glyoxylate and dicarboxylate metabolism, and taurine and hypoxanthine metabolism (Fig 7A; Supplementary Figs 8-11). Iron metabolism genes, too, differed between wild-type HID and the other dauer larvae, but gene expression was significantly and strongly correlated amongst these other dauer larvae. In other pathways too, such molybdenum cofactor biosynthesis pathway and PUFA biosynthesis, the HID differed from all dauer except *daf-2* dauers, and in chitin degradation and galactose metabolism, they differed from all other dauer except *cep-1* oe dauers, with whom they showed a significant and high correlation of gene expression pattern (Fig 7A; Supplementary Figs 8-11).

**Figure 7.**
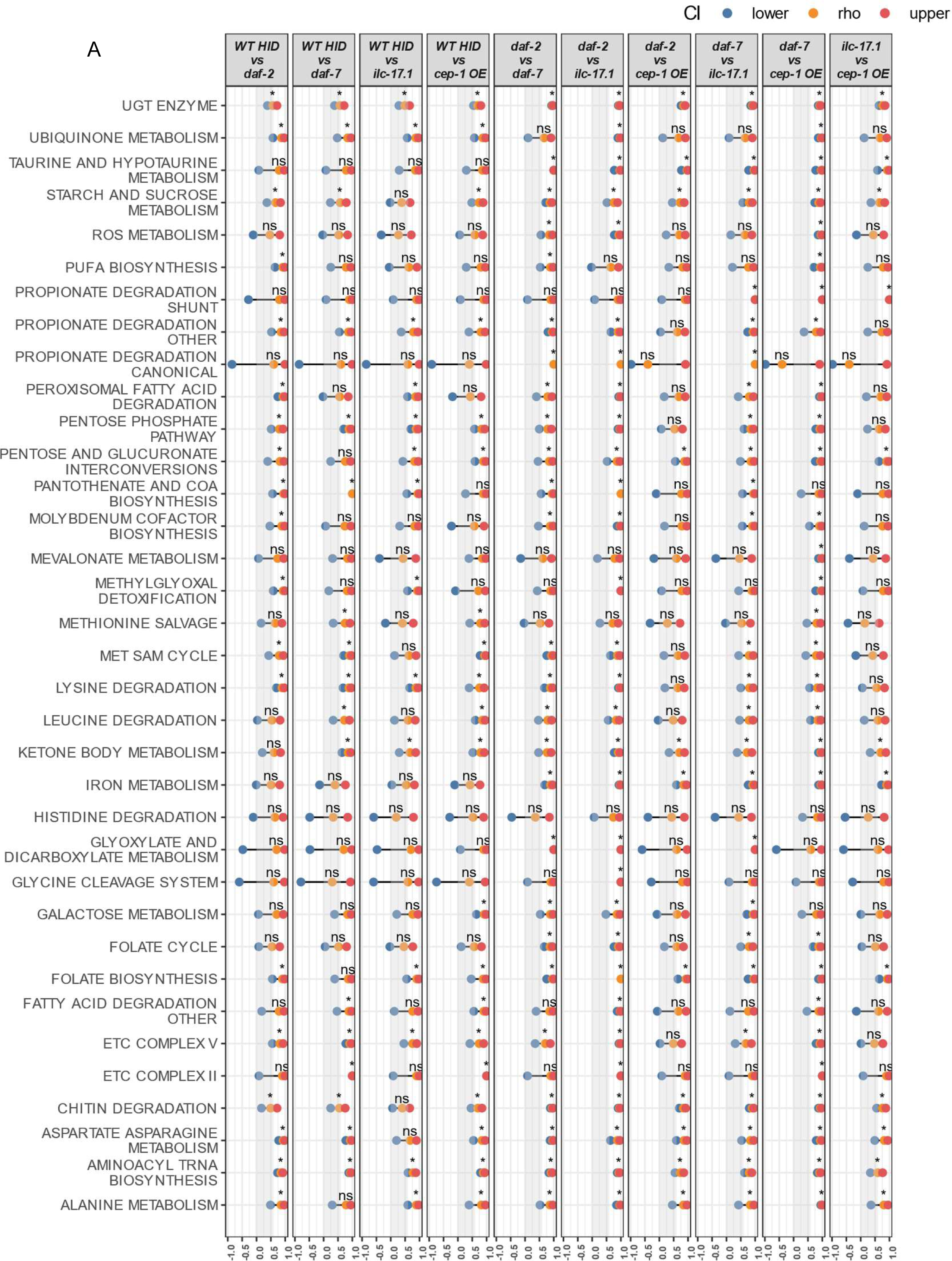
Correlation between dauer larvae in metabolic pathway-gene expression reveals broad variation. **A.** Dumbbell Plot showing Spearman’s correlation coefficient (rho) between pairs of dauers compared within metabolic pathways identified in the iCEL1314 metabolic network [Nanda et al. (2023)]. TOP labels: the pair of dauers compared. Y-axis: pathway. X-axis: rho value and confidence interval (CI): blue dot represents lower CI, yellow dot represents rho value, and red dot depicts high CI. p-values, Bonferroni corrected. *p-value <0.05; ns, not-significant.

Similarly, *daf-2* and *ilc-17.1* dauers differed from all other dauers in the expression of genes in the methionine salvage pathway, while the remaining dauers, wild-type HID, *cep-1 oe* and *daf-7* showed a strong and significant correlation in the expression of these genes [rho=0.7-0.8, padj<0.05; (Fig 7A; Supplementary Tables 17 and 18). The expression of genes involved in fatty acid degradation, Complex II activity and glycine cleavage system were also different in *daf-2* and *ilc-17.1* dauers compared to all other dauers. There were also differences in gene expression patterns between only specific pairs of dauer larvae. For instance, *daf-2* dauers differed from wild-type HID dauers in their expression of genes involved in leucine degradation [padj=0.6, and not significant] and also differed from *cep-1 oe* dauer larvae in gene expression related to leucine degradation and molybdenum cofactor biosynthesis [padj=0.3 and 0.2, respectively, and not significant]. Wild-type HID dauers and *ilc-17.1* dauers differed in the expression of genes involved in the chitin degradation pathway [rho=0.4, padj<0.05], leucine degradation pathway [rho=0.6, padj=0.1 and not significant], molybdenum cofactor biosynthesis [padj=0.1 and not significant], and starch and sucrose metabolism pathways [padj=0.1 and not significant]. The latter difference supported previous observations that *ilc-17.1* larvae might be deficient in their ability to take up glucose from their dietary source. Dauers induced by the loss of *daf-7* function showed the least differences; they too differed from wild-type HID dauers in the expression of genes in the molybdenum cofactor biosynthesis pathway [padj=0.1 and not significant], but otherwise largely shared gene expression patterns with more than one other type of dauer larva. Gene expression in the Met/SAM cycle differed between *cep-1 oe* on the one hand and *ilc-17.1* and *daf-2* on the other (Fig 7A; Supplementary Tables 17 and 18). We summarize all the differences between the dauers (Figure 8).

**Figure 8.**
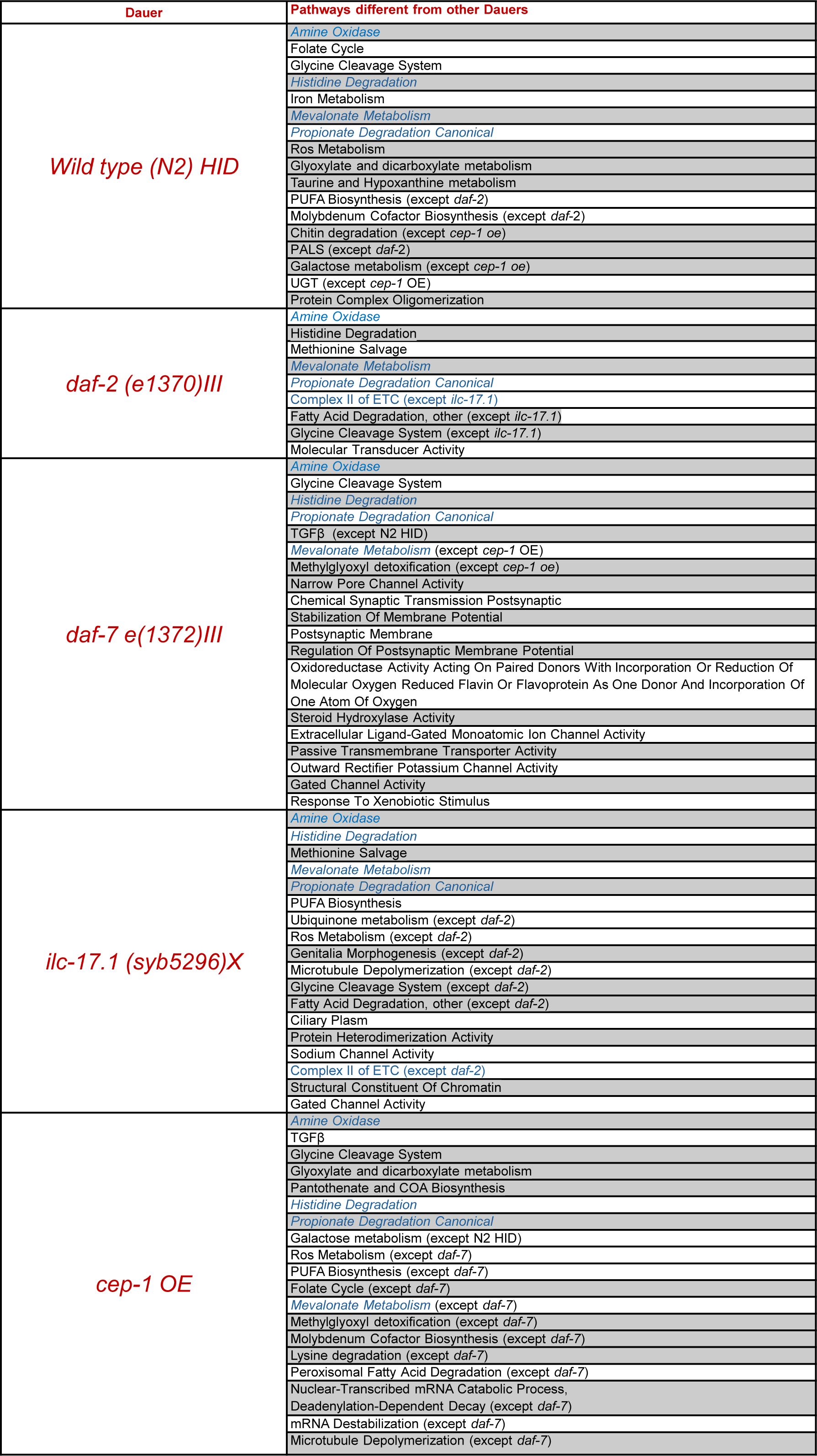
Gene expression differences between dauer larvae induced to arrest development by different stimuli. A limited number of gene expression pathways that regulate specific morphological features, and specific stress, immune, detoxification and metabolic programs vary between the different dauer larvae. These are summarized here.

This pervasive variability in the metabolic pathways was also confirmed upon comparing the correlation between genes in all metabolic pathways with those in all the Kim mountains, where except between wild-type HID dauers and *cep-1 oe* dauers, the metabolic pathway genes were significantly lesser correlated between all dauers (Fisher r-to-z transformation to calculate the significance of difference between rhos; z=-10.78 to 3.98 in favor of the Kim mountains, padj. <0.05; Supplementary Table 19).

## DISCUSSION

In summary, we show that the expression of individual genes in different *C. elegans* dauer larvae vary, as indicated by the fact that most genes have >50% CVs. The high CVs likely result from the similarities and differences in the expression of different subsets of genes in different subsets of dauers. Notwithstanding this high variability, with the exception of a limited number of pathways prominently involved in stress, metabolism, and the regulation of specific morphological traits, gene expression patterns between the different dauer larvae are strongly correlated, suggestive of transcriptional robustness. We speculate that during dauer entry, as seen during other developmental programs, there are gene regulatory mechanisms that stabilize gene expression variation to generate functional outcomes, which, in this case, is the developmental arrest of larvae in a hypometabolic dauer state. Previous studies have shown that the gene regulatory networks and other contexts within which genes operate constrain their variance and gene expression patterns ^1,2,5,6,32,33,35,45^. Our analysis supports this conclusion, showing that gene expression in the core dauer pathway involved in the dauer decision and arrest, i.e., the ILS (insulins and DAF-16 targets), TGF-β, DAF-12/steroid hormone pathways, cell cycle regulation, glycolysis, and ETCs pathways is robust and strongly correlated between dauer larvae irrespective of the stimulus that triggered dauer entry. Expression of cuticulins varies both at the level of expression as well as in its correlation; however, whether the alae structure itself varies between dauers was not examined.

The mechanisms and functional consequences of the observed variation in gene expression in pathways that regulate morphological features such as dendrite arborization and specific stress, immune, detoxification, and metabolic programs are unclear. However, the existence of this variability suggests that dauers generated by different triggers possess different suites of traits that may allow for better adaptation to specific niches. For instance, the route of dauer entry could influence the dramatic remodeling of neuron morphology and dendrite arborization^46–49^ particularly in the IL2 neurons, which are responsible for the dauer-specific behaviors, and the known sex bias in the ability of larvae to survive dauer^50,51^. The most pervasive variation between the different dauers was seen in the expression of genes in metabolic pathways. The reason why metabolic enzymes are more variable in some dauer larvae is not clear. Some of these enzymatic pathways may be inherently more variable and thus may also vary between dauers. For instance, propionate metabolism has undergone changes in the nematode clade, varying between *C. elegans* and wild strains and being lost in several parasitic helminth species^52–55^. In addition, since dauers do not feed, they must metabolize stored macronutrients, mainly lipids, but also glycogen and trehalose, and the variability in specific metabolic pathways might be related to the food intake of these larvae prior to dauer arrest. Thus, the methionine salvage pathway, leucine degradation, and histidine degradation are related to the generation of the essential amino acids histidine, methionine, and leucine, which in *C. elegans*, are typically obtained from the microbial diet^52–55^. Similarly, moco enzymes that are central to the Molybdenum cofactor (Moco) biosynthesis pathway can be obtained from the organism’s microbial diet, or can be synthesized by the animal, and thus food intake and molybdate biosynthesis by the bacteria prior to dauer arrest might influence the expression patterns of Molybdenum cofactor (Moco) biosynthesis pathway genes^56,57^. However, it is also possible that these metabolic pathways act downstream of sensors that must exist in all dauer larvae to continually monitor the environment and control their entry or exit from dauer arrest. Indeed, of the pathways that varied, such a role has been reported for peroxisomal fatty acid degradation, where deficiency in peroxisomal fatty acid β-oxidation in ASK neurons leads to the premature interruption of the dauer arrest and promotes an untimely exit from dauer to promote continued development even under dauer-inducing conditions such as increased levels of ascarosides^58,59^. Such a role can be envisioned for metabolic pathways modulated by microbial metabolites, whose presence could signify the advent of favorable conditions.

Nematodes fill all trophic levels in the food web ^60^ and are the most abundant metazoan species on the planet ^61–63^ . Yet, the mechanisms that facilitate their adaptation to diverse and often hostile niches remain poorly understood. Almost all nematode species have evolved forms of hypobiosis to adapt their life cycles to variable, unpredictable, and harsh environmental conditions ^60–67^. , of which a specialized form is the developmental diapause ‘dauer’. Indeed, it has been hypothesized that among parasitic nematode species, the dauer stage may have been a prerequisite for the evolution of the diversity of their parasitic lifestyles^60–67^ Thus, it is tempting to speculate that the transcriptional robustness of core dauer pathways allows for buffering and, thus, variation in the expression of genes involved in their response to the environment. Such differences could be pivotal in allowing the different dauers to be better suited to survive in and colonize diverse niches.

## Acknowledgements

We thank the V.P. laboratory and Dr. Gidalevitz (Drexel University) and Dr. He (University of Iowa) for their comments. Nematode strains were provided by the Caenorhabditis Genetics Center (CGC) (funded by the NIH Infrastructure Programs P40 OD010440). This work was supported by NIH R01 AG060616 (V.P.) and by National Cancer Institute (NCI) grant P30CA016056 involving the use of Roswell Park Comprehensive Cancer Center’s Pathology Network, Genomic, and Biomedical Research Informatics Shared Resources.

## Author Contributions

All authors designed the study and performed experiments. J.C.C. and V.P. designed the analyses, J.C.C. conducted the analysis data, and J.C.C. and V.P. drafted the manuscript.

## METHODS AND MATERIALS

### Growth conditions for *C. elegans* strains

All strains were grown and maintained at 20°C unless otherwise mentioned. Animals were grown in 20°C incubators (humidity controlled) on 60mm nematode growth media (NGM) plates by passaging 8-15 L4s (depending on the strain) onto a fresh plate. Extra care was taken to ensure equal worm densities across all strains. Animals were fed *Escherichia coli* OP50 obtained from Caenorhabditis Genetics Center (CGC) that were seeded (OD_600_=1.5 and this was strictly maintained throughout the experiments) onto culture plates 2 days before use. The NGM plate thickness was controlled by pouring 8.9ml of autoclaved liquid NGM per 60mm plate. Laboratory temperature was maintained at 20°C and monitored throughout. For all experiments, age-matched day-one hermaphrodites, or larvae timed to reach specific developmental stages as mentioned in the figure legend, were used.

#### C. elegans strains

*C. elegans* strains used in this study are listed in Table 1. Strains were procured from Caenorhabditis Genetics Center (CGC, Twin Cities, MN), generated in the laboratory or generated by Suny Biotech (Suzhou, Jiangsu, China 215028).

**Table 1.**

| Strain Name | Gene name | Source | Additional information |
| --- | --- | --- | --- |
| N2, *C. elegans* var *Bristol* | N2, Wild-type | Caenorhabditis Genetics Center |  |
| CB1370 | *daf-2* (e1370) III | Caenorhabditis Genetics Center |  |
| VEP032 | *ilc-17.1* (syb5296) X | Prahlad Lab/<br>SunyBiotech | *ilc-17.1* deletion, 2173bp deletion, and the 15bp and 127bp sequences were left in the 5' and 3' deletion end, respectively of the 2135 bp *ilc-17.1* gene |
| VEP036 | *unc-119* (ed4); *gtls1* [CEP-1::GFP + *unc-119* (+)] | Prahlad Lab | CEP-1 overexpression (ref: <sup>68</sup>) |
| CB1372 | *daf-7*(e1372) III. | Caenorhabditis Genetics Center |  |

### Obtaining larvae and dauers following ‘bleach-hatching’

Populations of 250-300 gravid adults were generated by passaging L4s on NGM plates. These plates were used for obtaining synchronized embryos by bleach-induced solubilization of the adults to then obtain larvae for harvesting mRNA. Specifically, animals were washed off the plates with 1X PBS and pelleted by centrifuging at 2665Xg for 30s. The PBS was removed carefully, and worms were gently vortexed in the presence of bleaching solution [250µl 1N NaOH, 200µl standard (regular) bleach and 550µl sterile water] until all the worm bodies had dissolved (approximately 5-6 minutes), and only eggs were viable. The eggs were pelleted by centrifugation (2665Xg for 45s), bleaching solution was carefully removed and then embryos were washed with sterile water 4-5 times and counted under the microscope. The desired number of embryos were seeded on fresh OP50 plates and allowed to grow at 25°C or 27°C for specific time periods depending on the experimental need. If >5% of eggs remained unhatched, these plates were discarded.

#### • 32 hrs L2/L3 larvae

Day-one adult worms grown at 20C (I-36LLVL Incubator) were bleach-hatched and ∼3200 eggs/genotype (∼800 eggs/plate and 4 plates/genotype) were seeded on fresh OP50 plates and allowed to grow for either 15 hours or 32 hours at 25°C in Echotherm incubator IN30-2. Worms were washed with sterile water, and total RNA was extracted from biological triplicates using the Direct-zol RNA Miniprep (catalog no. R2050, Zymo Research).

#### • 48 hrs dauer larvae

Day-one adult worms grown at 20C (I-36LLVL Incubator) were bleach-hatched, and ∼3200 eggs/genotype (∼800 eggs/plate and 4 plates/genotype) were seeded on fresh OP50 plates and allowed to grow for 48 hours at 25°C Echotherm incubator IN30-2. All strains other than N2 (wildtype control) showed >99.5% dauers at 48hrs. Any non-dauer worms were picked off the plate before collection to avoid variable staged worms in the RNA prep. N2 (wildtype control) dauers were obtained by incubating the eggs for 48 hours at 27°C in New Brunswick Galaxy 170S Incubator. These plates consisted of >96% dauers and all non-dauer worms were picked off to avoid variable staged worms in the RNA prep. Worms were washed with sterile water, and total RNA was extracted from biological triplicates using the Trizol extraction method.

### RNA extraction methods

#### Trizol extraction

300 µl of Trizol (catalog no. 400753, Life Technologies) was added to the samples after collection and snap-frozen immediately in liquid nitrogen. Samples were thawed on ice and then lysed using a Precellys 24 homogenizer (Bertin Corp.). RNA was then purified as detailed with appropriate volumes of reagents modified to 300 µl of Trizol. The RNA pellet was dissolved in 17 µl of RNase-free water. The purified RNA was then treated with deoxyribonuclease using the TURBO DNA-free kit (catalog no. AM1907, Life Technologies) as per the manufacturer’s protocol. cDNA was generated by using the iScript cDNA Synthesis Kit (catalog no. 170–8891, Bio-Rad). qRT-PCR was performed using PowerUp SYBR Green Master Mix (catalog no. A25742, Thermo Fisher Scientific) in QuantStudio 3 Real-Time PCR System (Thermo Fisher Scientific) at a 10 µl sample volume in a 96-well plate (catalog no. 4346907, Thermo Fisher Scientific). The relative amounts of mRNA were determined using the ΔΔCt method for quantitation. We selected pmp-3 as an appropriate internal control for gene expression analysis in C. elegans.

All relative changes of mRNA were normalized to either that of the wild-type control or the control for each genotype (specified in figure legends). Each experiment was repeated a minimum of three times. For qPCR reactions, the amplification of a single product with no primer dimers was confirmed by melt-curve analysis performed at the end of the reaction.

### Direct-zol RNA Miniprep (catalog no. R2050, Zymo Research)

Instructions on the kit were followed.

An equal volume of ethanol (95-100%) was added to the samples and mixed thoroughly. The mixture was transferred into a Zymo-Spin™ IICR Column in a Collection Tube and centrifuged at 10,000-16,000 x g for 30 seconds. The column was transferred into a new collection tube, and the flow-through was discarded. The sample was treated with DNase I and incubated at room temperature (20-30°C) for 15 minutes. 400 µl Direct-zol™ RNA PreWash was added to the column and centrifuged at 10,000-16,000 x g for 30 seconds. The flow-through was discarded, and the previous step was repeated. 700 µl RNA Wash Buffer was added to the column and centrifuge for 1 minute to ensure complete removal of the wash buffer. The column was transferred carefully into an RNase-free tube. RNA was eluted by adding 50 µl of DNase/RNase-Free Water directly to the column matrix and centrifuged at 10,000-16,000 x g for 30 seconds.

### RNA-seq analysis

RNA seq data for C. elegans dauer was analyzed and processed using nf-core/rnaseq v3.12.0^69^, and the pipeline was executed with Nextflow v22.10.6^70^. In short, the quality of the reads was assessed with Fastqc v0.11.9, and Trimgalore v0.67^71^ was used to filter low quality reads and remove adapters. The processed reads were to aligned to *C. elegans* genome (WBcel235 Ensembl release 111^72^), using STAR v2.7.9a^73^ with default settings. Alignments were quantified with Salmon using the WBCel235 annotation. Quality control of the alignment was performed with Qualimap v2.2.2^74^ and RseQC v3.0.1^75^.

#### • Differential Expression and Normalization

The transcript-level estimates were summarized to gene-level counts and to gene-level Transcript per million (TPM) with the tximport package^76^. The RNA-seq data from the L2/L3 larvae (WT at 32hours) was analyzed as previously described^77^. Gene-level counts from this study were merged with the Dauer gene-level counts to perform the normalization and differential expression steps. DESeq2^78^ was used to do differential expression between the samples (Supplementary Table 20). Genes with low read counts (n <10) were removed from the differential expression analysis. Genes with an adjusted p-value of <0.05 (after correction with Benjamini & Hochberg) were considered significant. Changes in expression were presented as log_2_ fold-change.

To obtain the mean gene expression, gene-level TPM, for each sample replicate (dauers only) were transformed log_10_ (TPM+1) and averaged between the replicates.

#### • Gene expression variability

Principal Component Analysis (PCA) and pairwise distance analysis were performed after transforming the raw counts with variance stabilization transformation (VST). PCA was done using the genes with highest variance (top 500), meanwhile the pairwise distances were calculated by determining the Euclidean distance between all the genes and then using hierarchical clustering. Regularized Log Transformation (rlog) transformation for Gene-level counts was used to determine gene variability and expression (Heatmap and meanSdPlot). Coefficient of Variation (CV) of the raw gene-level counts was used to determine the level of gene variability across all samples. Genes were grouped into clusters by calculating the Z-score of the gene expression (log_10_ TPM+1), then applying hierarchical clustering (distance: Euclidean, method: complete). The optimal number of clusters was determined by using the elbow method.

#### • Correlation Analysis

Spearman’s correlation and confidence intervals were calculated using the R package psych ^79^. Global gene correlation was calculated by filtering genes with a TPM value <1, and then plotting them as log_10_TPM values in scatter plots with the R package ggpubr^80^.

Correlations for Pathways or GO categories between the different types of dauers (WT HID, ilc-, daf-2, daf-7, cep-1 OE) were calculated by using the log_10_(TPM+1) values of the specified subset of genes, then plotted as dumbbells or heatmap, p-values were adjusted for multiple tests using Bonferroni correction. Rho values <0.6 were considered low, and adjusted p-values <0.05 were considered significant.

The significance of the difference between the correlation coefficients was done by performing a Fisher Z transformation with R package TOSTER^81^. P-values values <0.05 were considered significant.

#### • Functional Analysis

Gene Ontology Analysis was performed using ClusterProfiler^82^ and Wormbase Enrichment Suite^83^. Ontology terms were obtained from the R package org.Ce.eg.db^84^ and from Wormbase ^85^. DEGs and CV ranked genes were used to perform an Over Representation Analysis (ORA). Ontology terms with an adjusted p-value <0.05 (Benjamini & Hochberg) or q-value <0.05 were considered significant.

To summarize and visualize the Gene Ontology terms the package rrvgo^86^ was used. Briefly, the GO terms were grouped by semantic similarity and simplified to their parent terms, using a 90% similarity threshold. Enriched parent terms were plotted using a UMAP projection.

#### • Visualization

Venn diagram was generated using the package Venn^87^. Heatmaps were generated using the package ComplexHeatmap^88^ and pheatmap^89^. All the statistical analyses were done in R ^90^.

## Data Availability

The dauer expression data have been deposited in NCBI’s Gene Expression Omnibus ^91^ with the GEO accession number GSE274872. The WT L2/L3 (WT 32 hours) data was previously published in NCBI’s Gene Expression Omnibus with accession numbers GSE218596 and GSE229132

---

**Supplementary Figure 1.**
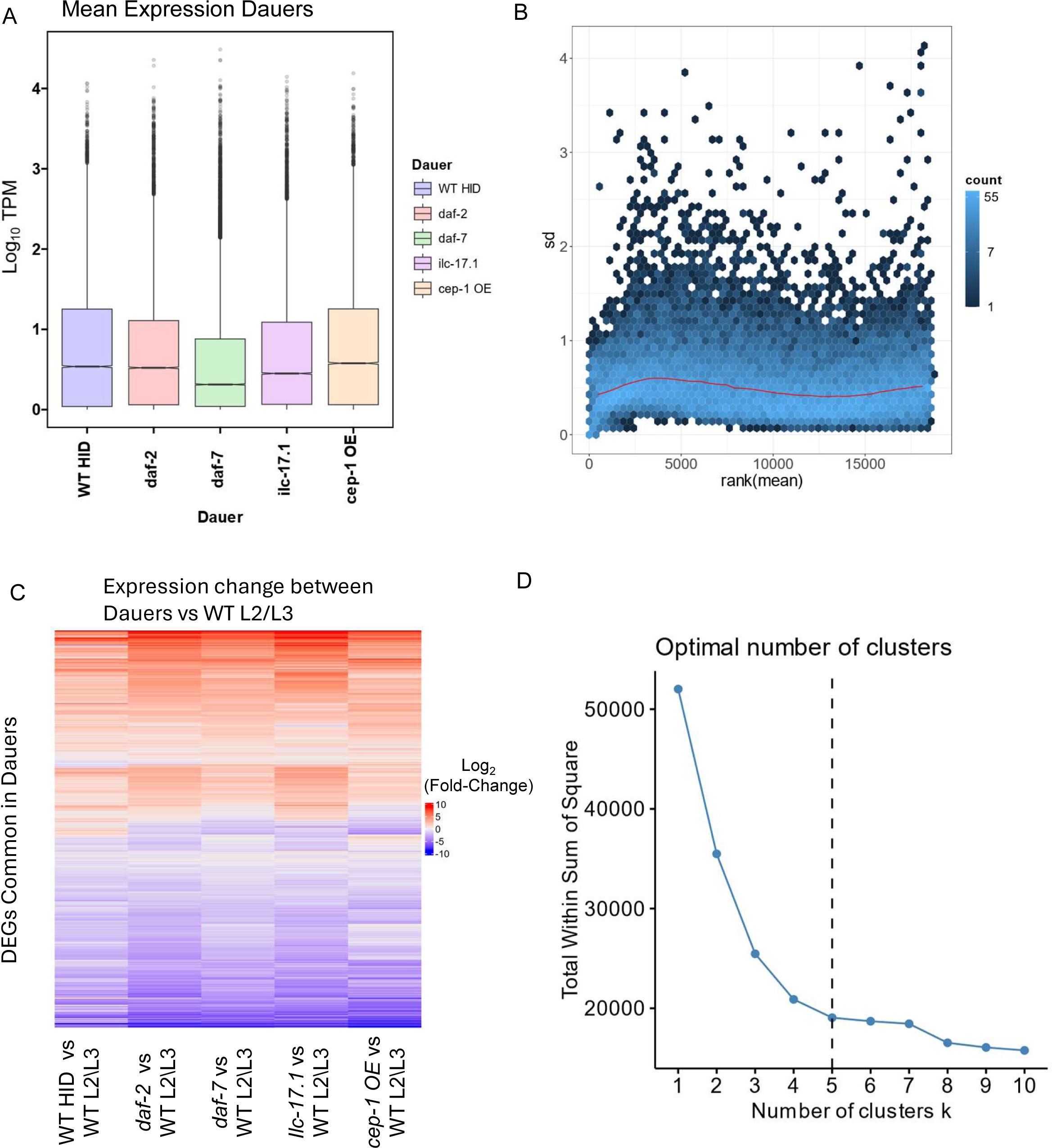
: Features of RNA-seq data. **A.** Boxplot showing the differences in gene expression in the different dauer larvae. X-axis: dauers [wild type HID (WT HID), *daf-2 (e1370) III*, *daf-7(e1372) III* and *ilc-17.1 (syb5296) X* and *cep-1 (cep-1oe)*]. Y-axis shows the Log10 (TPM+1) expression. **B.** Scatter Plot showing the Y-axis: Standard Deviation (SD) of the all genes, RLog transformed counts. X-axis: mean values (ranked) of all genes. **C.** Heatmap showing change in expression, Log_2_ Fold-Change, between dauers and wild-type L2/L3 in the 13120 DEGs shared by all dauers. **D.** Elbow Method showing significant drops in Within Sum of Square (WSS) around K = 5 (dotted line)

**Supplementary Figure 2:**
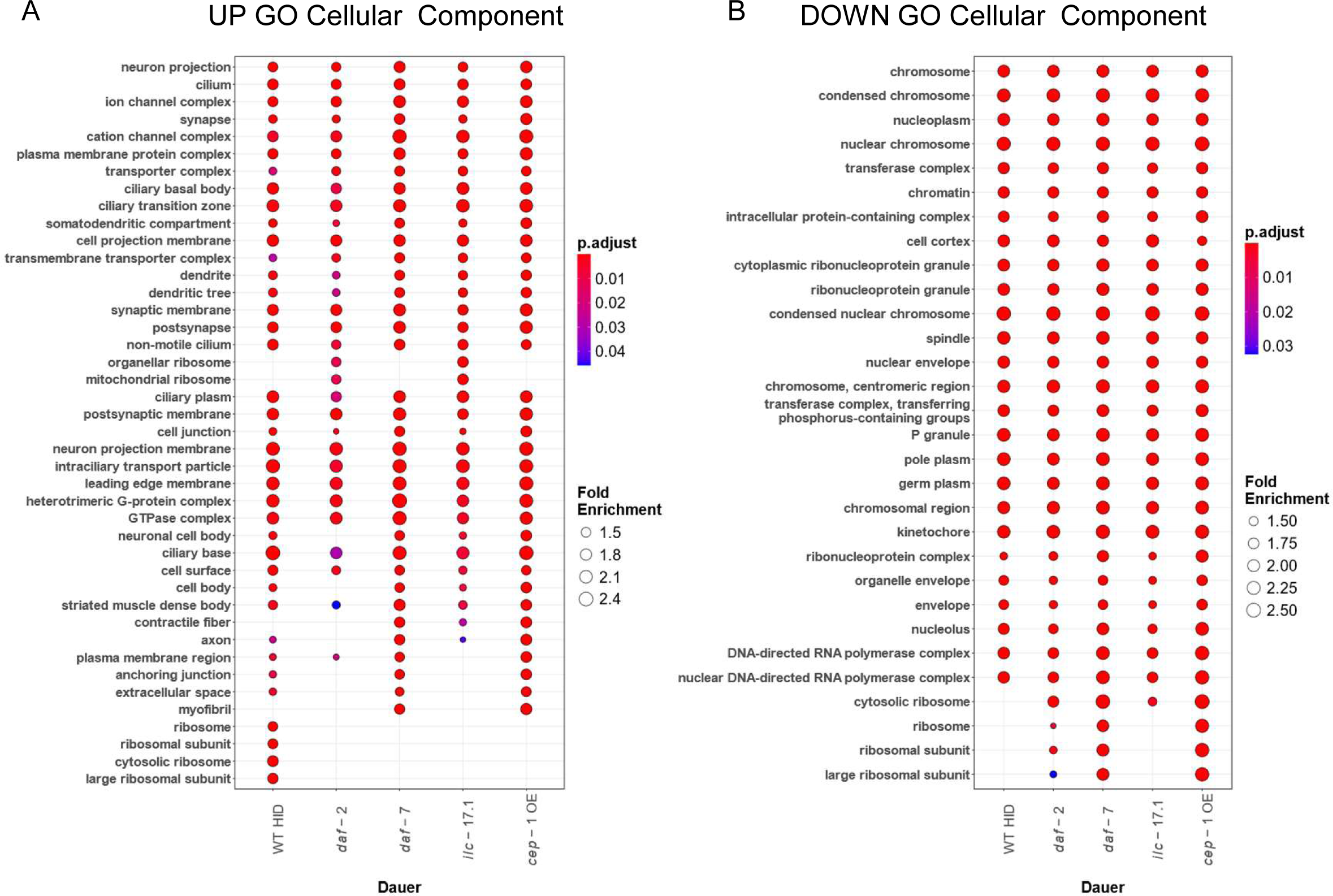
Gene Ontology Analysis: Pathways shared by all dauers, Cellular Component. **A.** Dot plot showing comparison of enrichment between upregulated DEGs in wild-type HID, *daf-2, daf-7*, *ilc-17.1 or cep-1oe* dauers. Y-axis : GO categories (Cellular Components). Color bar: adjusted p-values (Benjamini-Hochberg corrected, p<0.05), lower p-value in red, higher p-value blue. Circle size : Fold Enrichment (Gene Ratio/Background Ratio). **B.** Dot plot showing comparison of enrichment between downregulated DEGs in wild-type HID, *daf-2, daf-7*, *ilc-17.1 or cep-1oe* dauers. Y-axis : GO categories (Cellular Components). Color bar: adjusted p-values (Benjamini-Hochberg corrected, p<0.05), lower p-value in red, higher p-value blue. Circle size : Fold Enrichment (Gene Ratio/Background Ratio).

**Supplementary Figure 3:**
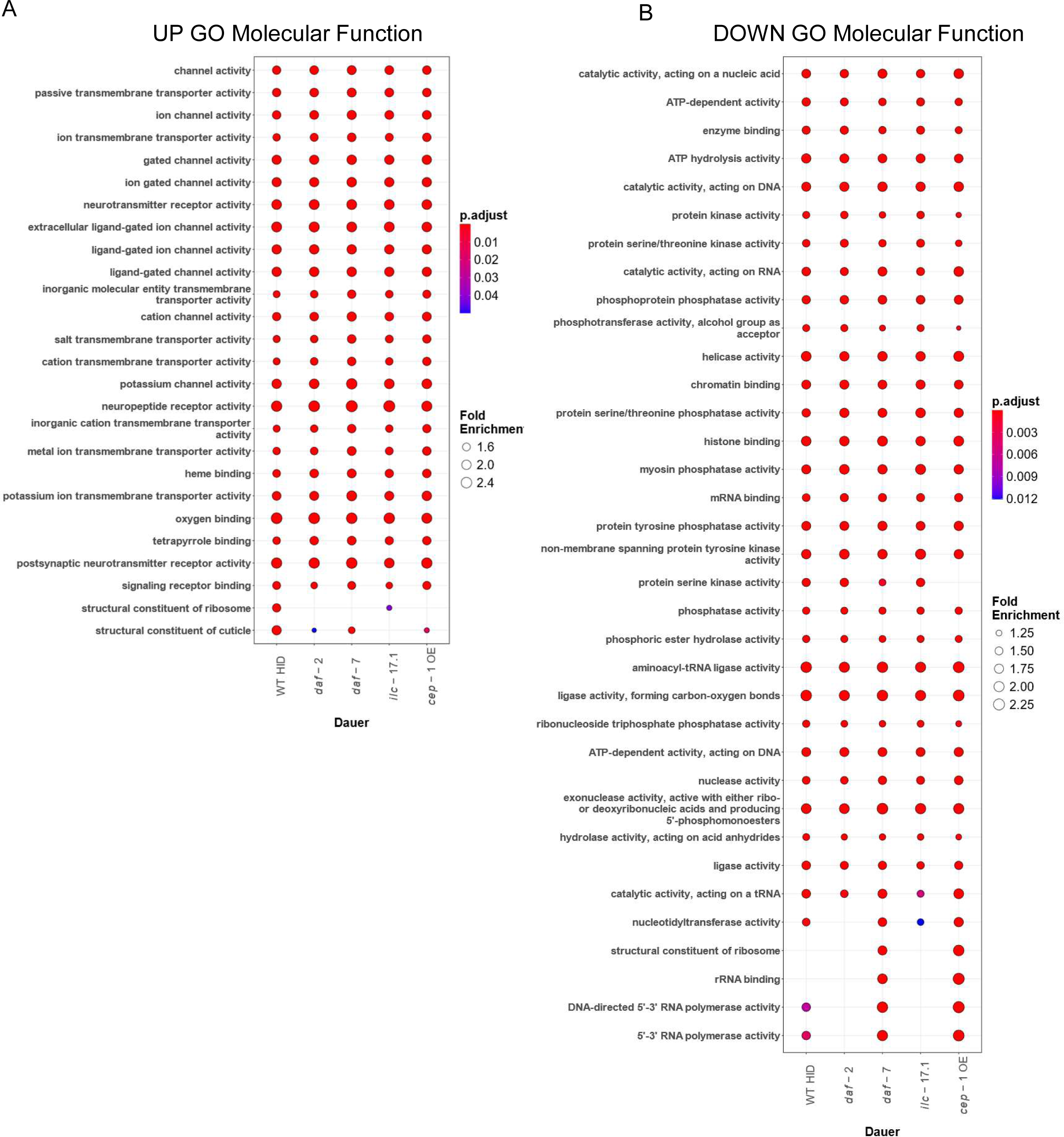
Gene Ontology Analysis: Pathways shared by all dauers, Molecular Function. **A.** Dot plot showing comparison of enrichment between upregulated DEGs in wild-type HID, *daf-2, daf-7*, *ilc-17.1 or cep-1oe* dauers. Y-axis : GO categories (Molecular Function). Color bar: adjusted p-values (Benjamini-Hochberg corrected, p<0.05), lower p-value in red, higher p-value blue. Circle size : Fold Enrichment (Gene Ratio/Background Ratio). **B.** Dot plot showing comparison of enrichment between downregulated DEGs in wild-type HID, *daf-2, daf-7*, *ilc-17.1 or cep-1oe* dauers. Y-axis : GO categories (Molecular Function). Color bar: adjusted p-values (Benjamini-Hochberg corrected, p<0.05), lower p-value in red, higher p-value blue. Circle size : Fold Enrichment (Gene Ratio/Background Ratio).

**Supplementary Figure 4:**
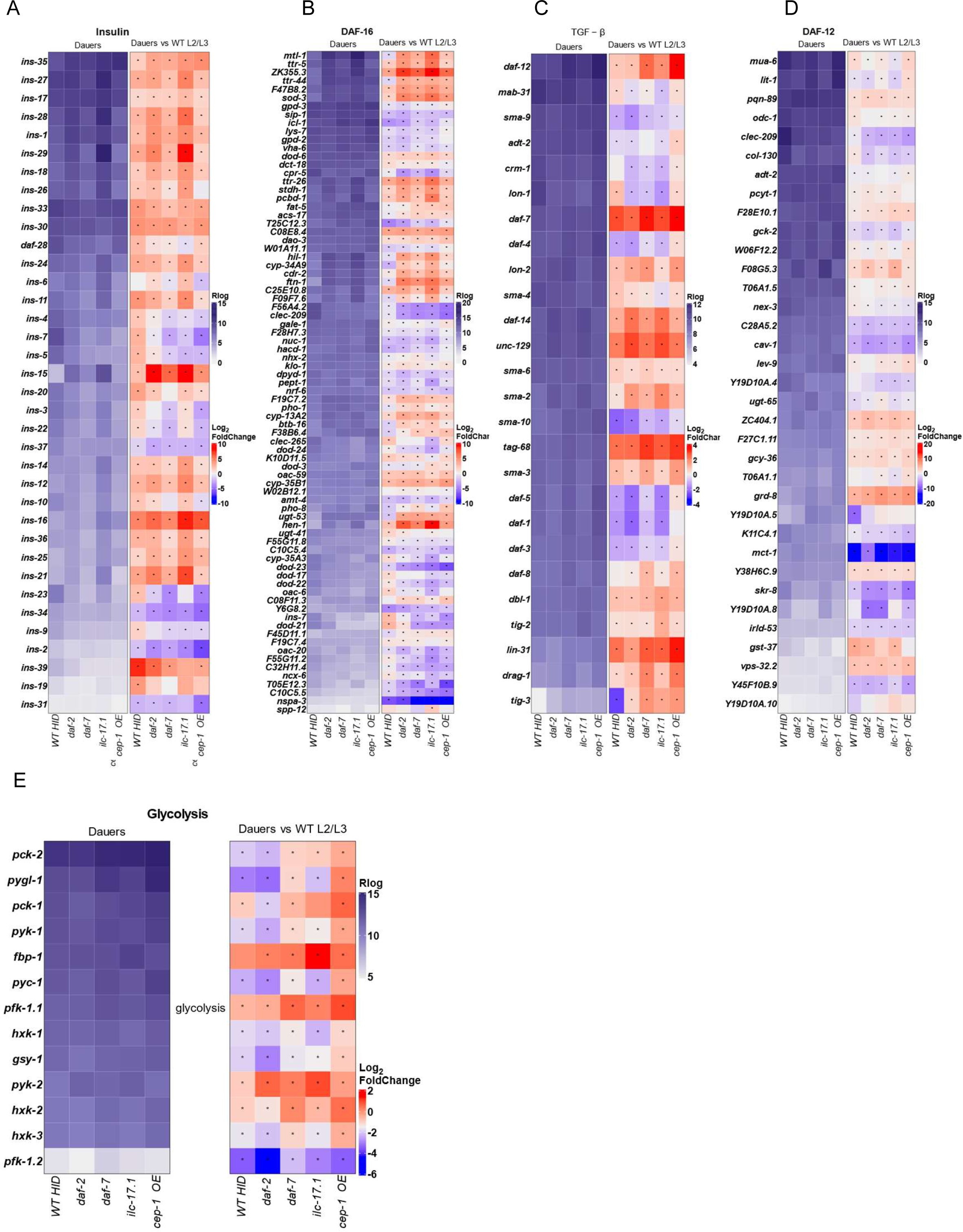
Heatmap of Gene Expression of the “core dauer pathway’ genes. **A-E**: Gene expression in A, Insulin pathway; B, DAF-16 pathway; C, TGF-β pathway; D, DAF-12 pathway and E, Glycolysis. Y-axis: gene names. **Left panel**: rlog transformed expression values for each dauer. Purple Color Bar: rlog expression. **Right panel**: Log_2_Fold change values, calculated for change between gene expression in dauer versus gene expression in wild-type L2/L3 larvae. Blue-Red color bar: Log_2_ Fold-Change (Dauer/larvae). Significant change: FDR<0.05. Bonferroni corrected.

**Supplementary Figure 5:**
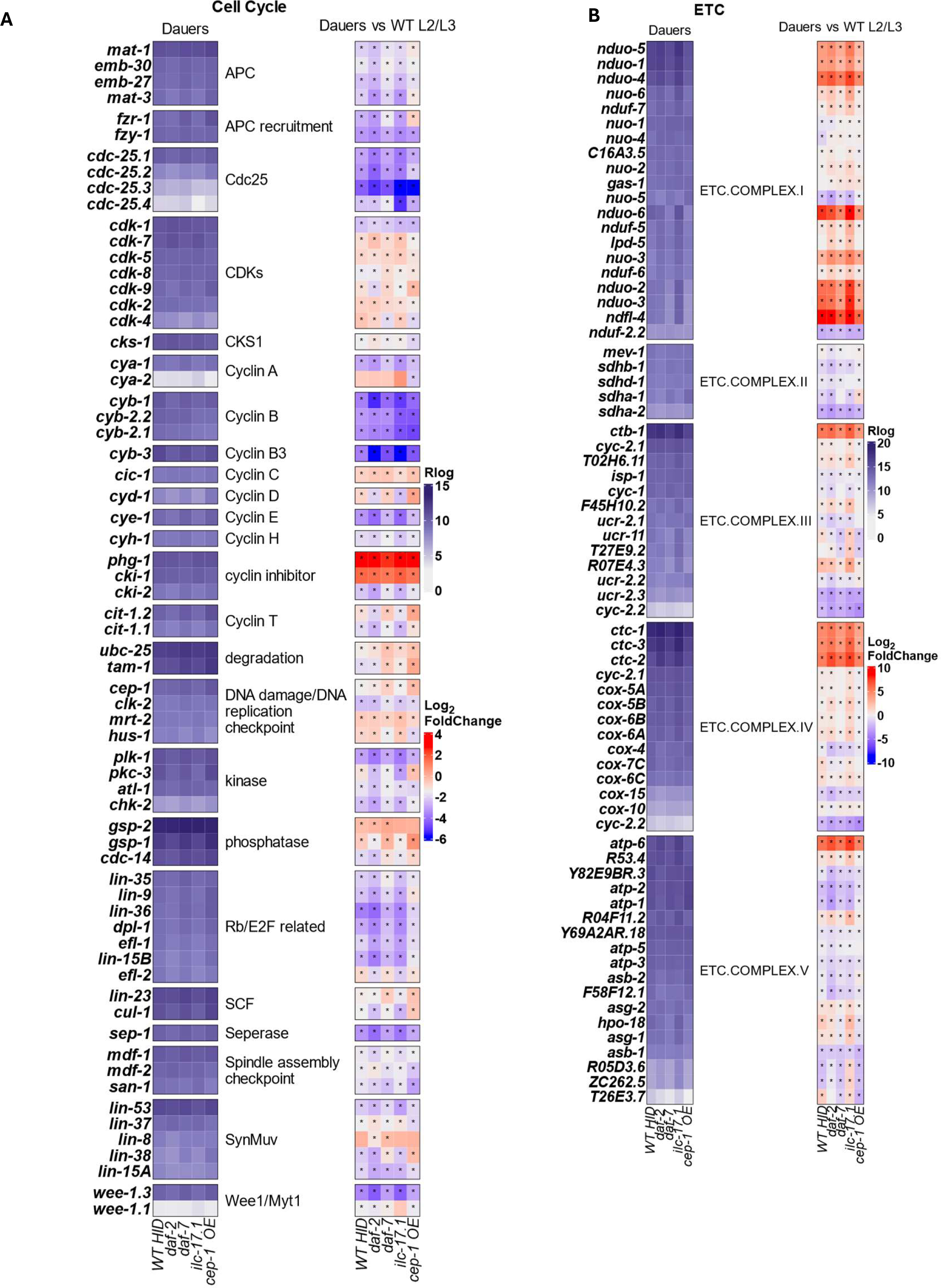
Heatmap of Gene Expression of the “core dauer pathway’ genes: cell cycle and electron transport chain (ETC) **(A-B)**: Gene expression in A, Cell Cycle genes; B, ETC components. Y-axis: gene names. **Left panel**: rlog transformed expression values for each dauer. Purple Color Bar: Rlog expression. **Right panel**: Log_2_Fold change values, calculated for change between gene expression in dauer versus gene expression in wild-type L2/L3 larvae. Blue-Red color bar: Log_2_ Fold-Change (Dauer/larvae). Significant change: FDR<0.05. Bonferroni corrected.

**Supplementary Figure 6:**
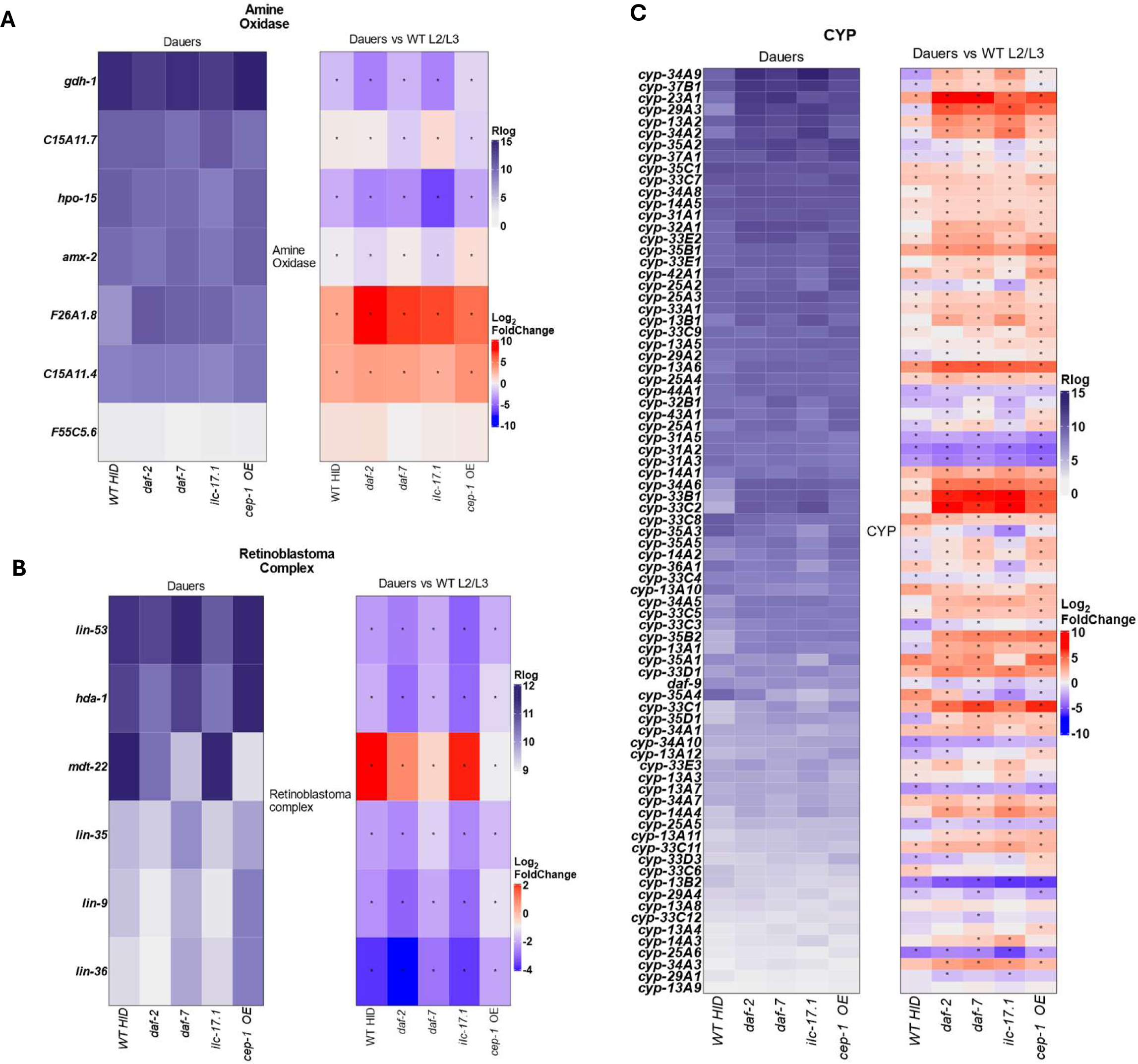
Heatmap of Gene Expression in Kim ‘mountains’ where the correlation between dauers is poor or not significant. **(A-C)** Gene expression in A, Amine Oxidases; B, Retinoblastoma components and C, Cytochrome P450 (CYP) genes. Y-axis: gene names. **Left panel**: rlog transformed expression values for each dauer. Purple Color Bar: rlog expression. **Right panel**: Log_2_Fold change values, calculated for change between gene expression in dauer versus gene expression in wild-type L2/L3 larvae. Blue-Red color bar: Log_2_ Fold-Change (Dauer/larvae). Significant change: FDR<0.05. Bonferroni corrected.

**Supplementary Figure 7:**
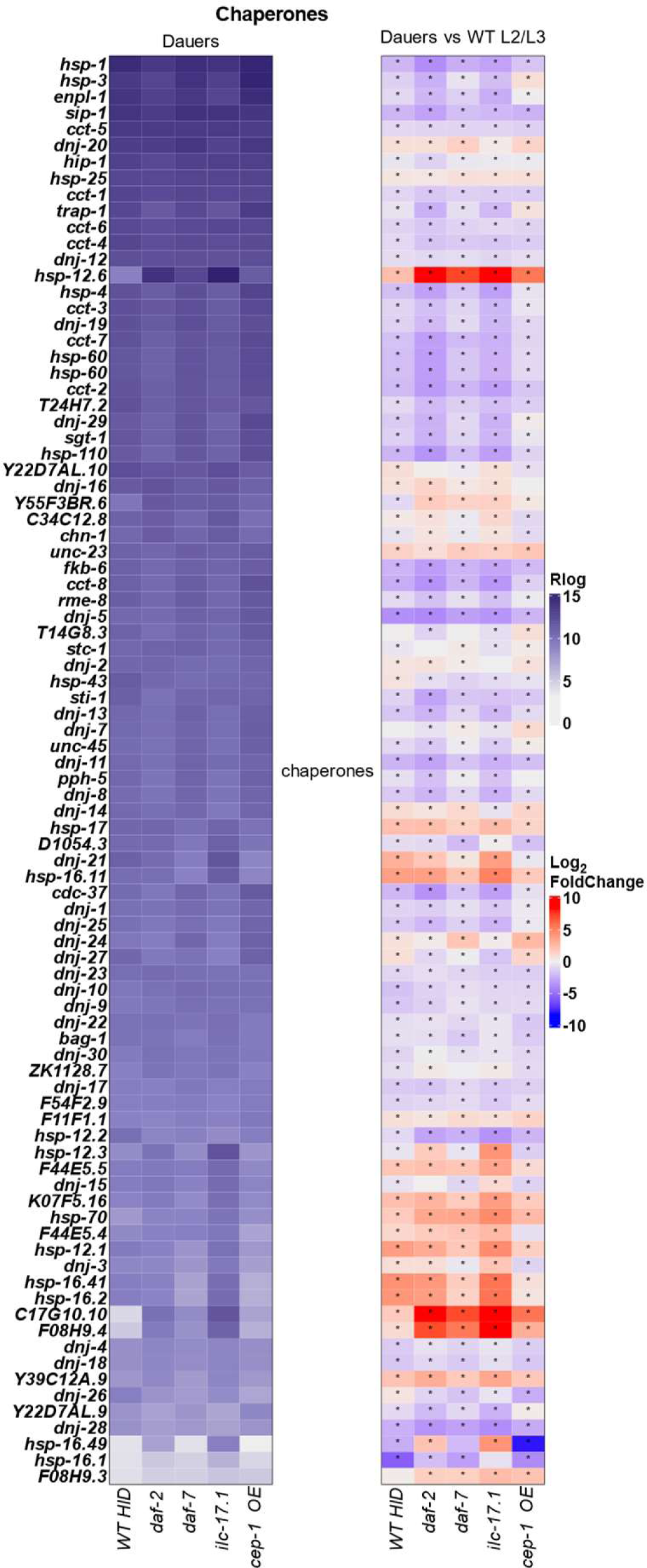
Heatmap of Gene Expression in Kim ‘mountains’: Heat shock (chaperones). Expression of chaperone genes. Y-axis: gene names. **Left panel**: rlog transformed expression values for each dauer. Purple Color Bar: rlog expression. **Right panel**: Log_2_Fold change values, calculated for change between gene expression in dauer versus gene expression in wild-type L2/L3 larvae. Blue-Red color bar: Log_2_ Fold-Change (Dauer/larvae). Significant change: FDR<0.05. Bonferroni corrected.

**Supplementary Figure 8:**
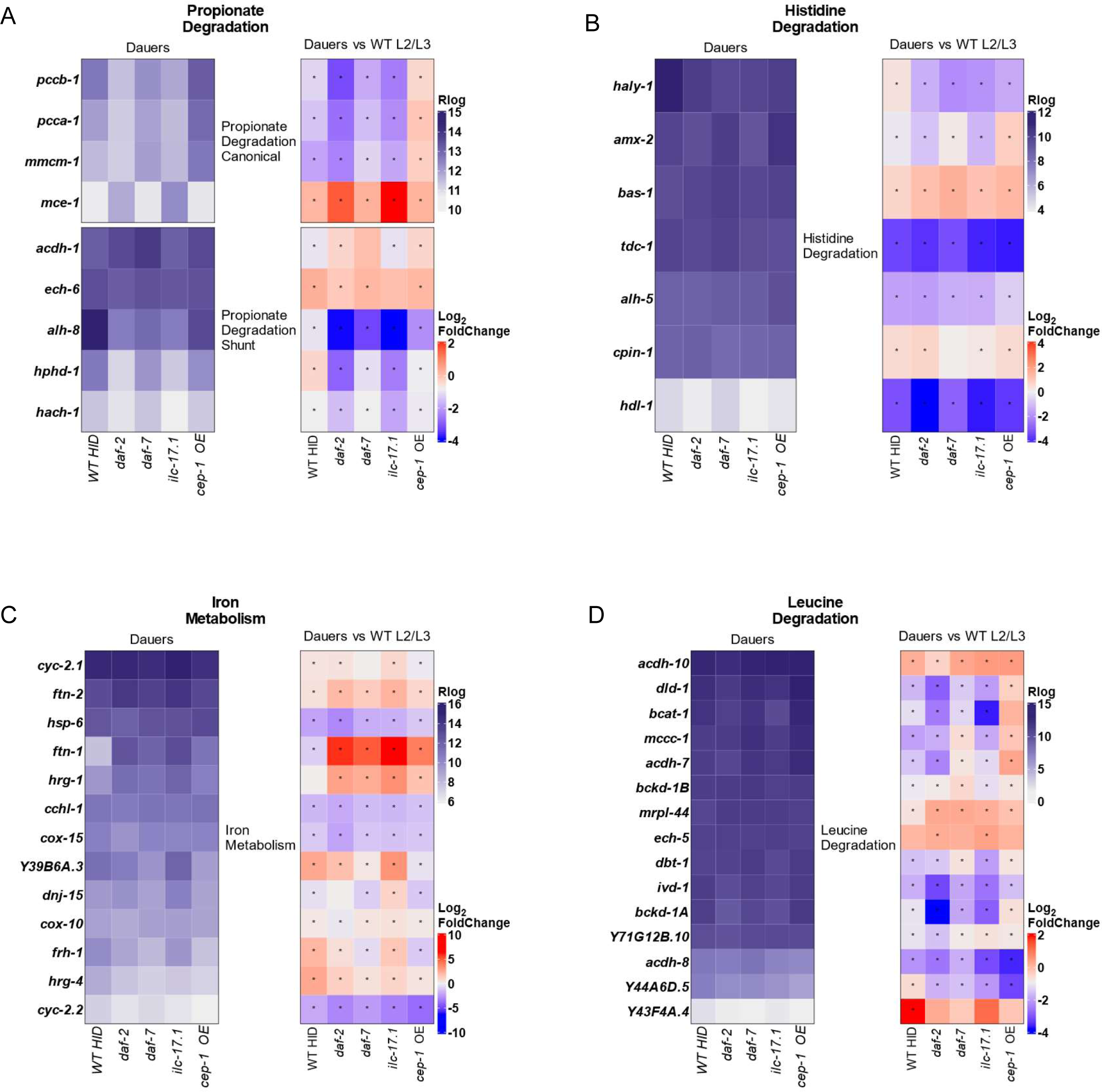
Heatmap of Gene Expression in Metabolic Pathways that vary between dauer larvae. **(A-D)** Gene expression in A, Propionate Degradation; B, Histidine degradation; C, Iron metabolism, and D, Leucine Degradation. Y-axis: gene names. **Left panel**: rlog transformed expression values for each dauer. Purple Color Bar: rlog expression. **Right panel**: Log_2_Fold change values, calculated for change between gene expression in dauer versus gene expression in wild-type L2/L3 larvae. Blue-Red color bar: Log_2_ Fold-Change (Dauer/larvae). Significant change: FDR<0.05. Bonferroni corrected.

**Supplementary Figure 9:**
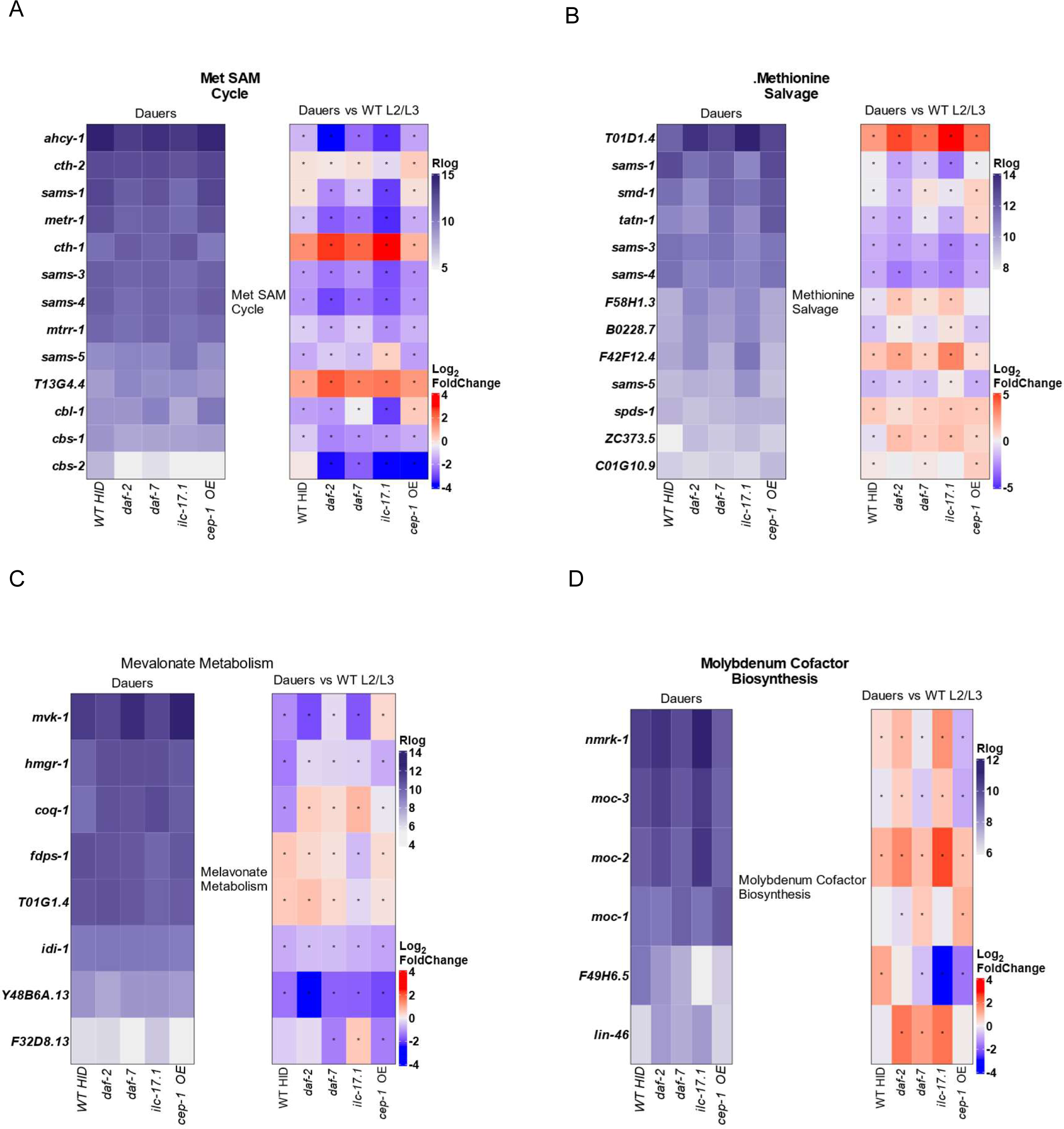
Heatmap of Gene Expression in Metabolic Pathways that vary between dauer larvae. **(A-D)** Gene expression in A, Met Sam pathway; B, Methionine salvage pathway; C, Mevalonate metabolism, and D, Molybdenum Cofactor Biosynthesis. Y-axis: gene names. **Left panel**: rlog transformed expression values for each dauer. Purple Color Bar: rlog expression. **Right panel**: Log_2_Fold change values, calculated for change between gene expression in dauer versus gene expression in wild-type L2/L3 larvae. Blue-Red color bar: Log_2_ Fold-Change (Dauer/larvae). Significant change: FDR<0.05. Bonferroni corrected.

**Supplementary Figure 10:**
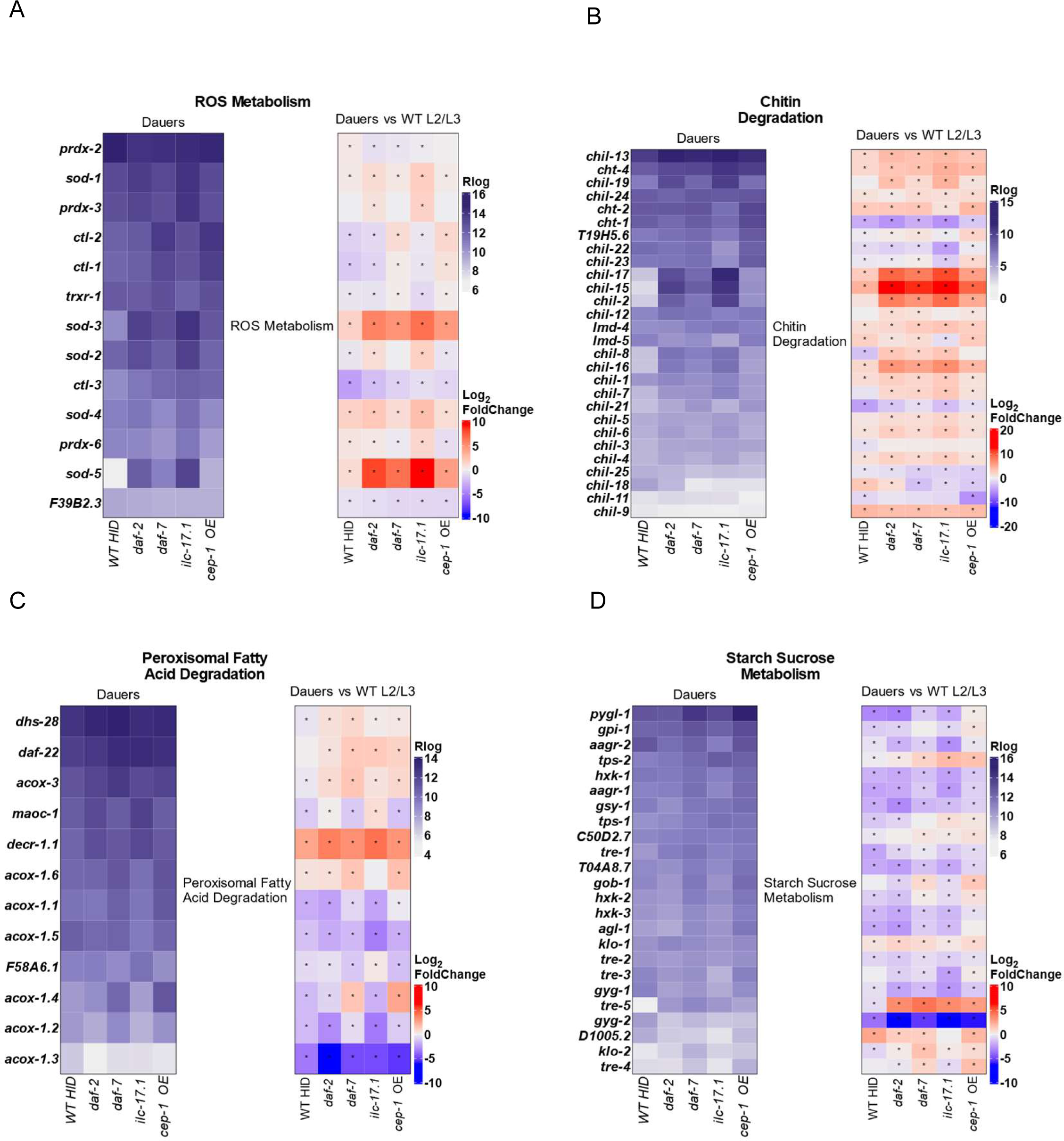
Heatmap of Gene Expression in Metabolic Pathways that vary between dauer larvae. **(A-D)** Gene expression in A, ROS metabolism; B, Chitin Degradation; C, Peroxisomal fatty Acid Degradation, and D, Starch, Sucrose metabolism. Y-axis: gene names. **Left panel**: rlog transformed expression values for each dauer. Purple Color Bar: rlog expression. **Right panel**: Log_2_Fold change values, calculated for change between gene expression in dauer versus gene expression in wild-type L2/L3 larvae. Blue-Red color bar: Log_2_ Fold-Change (Dauer/larvae). Significant change: FDR<0.05. Bonferroni corrected.

**Supplementary Figure 11:**
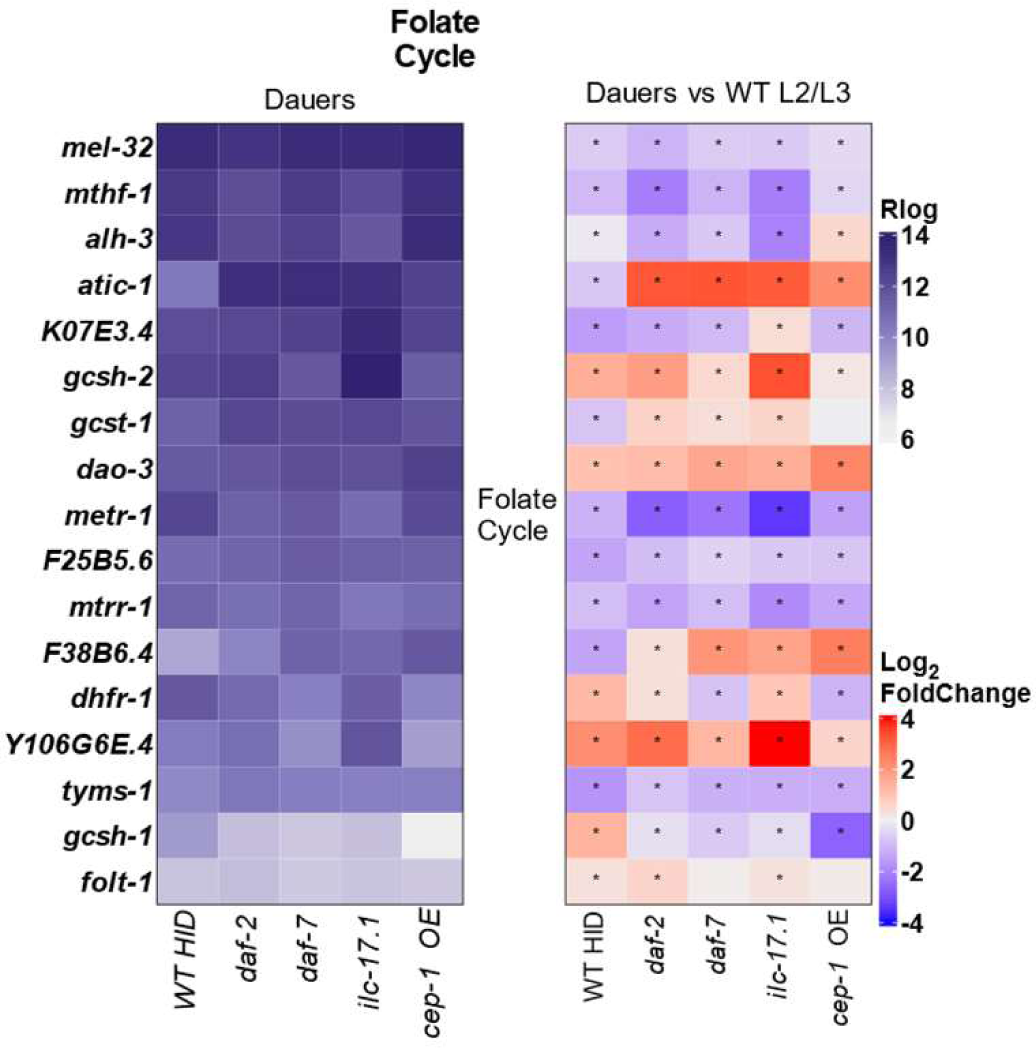
Heatmap of Gene Expression in the Folate Pathway varies between dauer larvae. Gene expression in the Folate Pathway. Y-axis: gene names. **Left panel**: rlog transformed expression values for each dauer. Purple Color Bar: rlog expression. **Right panel**: Log_2_Fold change values, calculated for change between gene expression in dauer versus gene expression in wild-type L2/L3 larvae. Blue-Red color bar: Log_2_ Fold-Change (Dauer/larvae). Significant change: FDR<0.05. Bonferroni corrected.

